# Lipid Network Crosslinked Hydrogels: Controlling Material Dynamics Across Multiple Length Scales Through Lipid Movement

**DOI:** 10.64898/2026.06.24.734376

**Authors:** Neil J. Baugh, Michelle S. Huang, Narelli de Paiva Narciso, Jordan A. Bunch, Jayniana Williams, Yueming Liu, Ruby Onsongo, David Kilian, Renato S. Navarro, Sarah C. Heilshorn

## Abstract

Control over network dynamics at different length scales is a feature of natural materials challenging to replicate in synthetic hydrogels. Hydrogel viscoelasticity is commonly controlled by tuning the kinetics of reversible crosslinks; however, this strategy inherently links the resulting macroscale and nanoscale dynamics of the individual network components. Taking inspiration from biological materials that feature lipids as structural elements, we introduce Lipid Network Crosslinked (LINC) hydrogels that exploit the mobility of individual lipids within self-assembled liposomes as covalent, network-crosslinking points. These mobile, covalent crosslinks increase hydrogel stress relaxation rates over 20-fold compared to polymer-only hydrogels with equivalent crosslinking chemistries and stiffnesses. We demonstrate that liposome design parameters, including degree of surface functionalization and tail saturation, provide a means to independently control the macroscale storage moduli and stress relaxation behavior. Finally, as an application where control over network dynamics at different length scales is critical, we placed cell-adhesive ligands onto more mobile or less mobile network elements. Human neural progenitor cells cultured within LINC hydrogels of identical macroscale viscoelasticity significantly altered their phenotype in response to nanoscale ligand dynamics. These results establish LINC hydrogels as biomimetic materials that leverage nanoscale lipid mobility within a macroscale polymeric network to control dynamics at multiple length scales.

## 1. Introduction

Native tissues have distinct time scales of motion at different length scales due to their hierarchical network structure that includes both lipids and polymers. At the macroscale, the extracellular matrix exhibits viscoelastic mechanical properties ^[1–3]^. At the nanoscale, the dynamics are dictated by the molecular mobility of individual network components, including the lipids within the cell membrane [4,5]. Recapitulating these multi-length scale dynamics in engineered materials remains a major challenge, as it is difficult to independently tune macroscale viscoelasticity and nanoscale mobility.

Viscoelastic hydrogels have a broad range of healthcare related applications, including as scaffolds for tissue engineering, delivery devices for regenerative medicine, and extrudable inks for 3D printing ^[6–10]^. Conventional viscoelastic hydrogels are formed by a network of polymers with reversible crosslinks, such as dynamic covalent chemistry (DCC) bonds or non-covalent interactions ^[6,11–13]^. These materials enable tuning of macroscale dynamics through control of molecular-level properties, including crosslink density and bond dynamics. In these systems, the macroscale stress relaxation rate is governed by the crosslink kinetics, which dictates nanoscale polymer mobility as the crosslinks reversibly detach and reform [14,15]. Alternatively, supramolecular assemblies can also form viscoelastic structural hydrogels [7,11,16,17]. Although nanoscale dynamics in supramolecular gels can be tuned through careful design, because these systems typically form larger length scale networks through entanglements of the individual matrix components, they lack independent control over macroscale dynamics ^[17–19]^.

Taking inspiration from natural tissues, we sought to leverage the mobility of lipids to design hydrogels with control over both macroscale and nanoscale dynamics. Lipids retain their nanoscale mobility upon self-assembly into larger structures [4,5,20], providing an attractive starting point to generate materials with tunable, multi-length scale dynamics. Lipid self-assembly is ubiquitous in naturally evolved systems, and while lipid-based particles are commonly used in hydrogel drug delivery ^[21–24]^, they have been overlooked as structural elements in hydrogel design. In certain conditions, lipids will spontaneously self-assemble into large lipid-rich domains in aqueous solvents [25], but do not form hydrogels without external bonding. At high concentrations, self-assembled lipids can form supramolecular hydrogels when combined with polymers containing hydrophobic pendants [26], fatty acids [27], nucleotide modifications [28], or high ion concentrations [29,30]. However, due to weak hydrophobic interactions these materials have limited mechanical properties and do not have multi-lengthscale dynamic control. Here, we introduce a novel structural hydrogel composed of nanoscale, self-assembled lipids that are covalently crosslinked into larger, macroscale, polymeric networks. This hierarchical design offers multi-length scale control of network dynamics, resulting in hydrogels with tunable macroscale viscoelasticity and nanoscale mobility.

Specifically, we design a supramolecular, Lipid Network Crosslinked (LINC) hydrogel, in which liposomes are covalently crosslinked to hyaluronic acid (HA) polymers through individual, mobile lipids (**Figure 1A**). This design allows for the presentation of lipid-tethered active elements (*e.g.* crosslinking ligands or cell-adhesive ligands) with tunable control over nanoscale dynamics in the mobile lipid bilayer (**Figure 1B**). We hypothesized that crosslinking the polymeric network through individual lipid-ligands would facilitate control over macroscale network dynamics, including stress relaxation behavior (**Figure 1C**). This design strategy exploits the structure-function relationships between lipid physicochemical properties and self-assembled membrane fluidity, providing a unique method to control hydrogel properties at both the nano- and macroscales. To validate this unique design strategy, we use fluorescence measurements of nanoscale lipid mobility and relate them to macroscale measurements of network rheology. This modular design flexibility enabled the formulation of hydrogels with a wide range of stiffnesses (shear moduli *G’* spanning about 100 to 2,000 Pa) and stress relaxation rates (relaxation half-lives spanning about 30 minutes to 24 hours). Importantly, we find that mobile crosslinking ligands increase hydrogel stress relaxation rates over 20-fold compared to polymer-only hydrogels with equivalent crosslinking chemistries and stiffnesses. Finally, we demonstrate that control over network dynamics at different length scales is critical in engineering cell morphology. By placing cell-adhesive ligands onto the more mobile lipid bilayers or less mobile polymer network, we show that human neural progenitor cells (hNPCs) cultured within LINC hydrogels of identical macroscale viscoelasticity significantly altered their morphology in response to nanoscale ligand dynamics. These results establish LINC hydrogels as biomimetic materials that leverage nanoscale lipid mobility within a macroscale polymeric network to control dynamics at multiple length scales.

**Figure 1.**
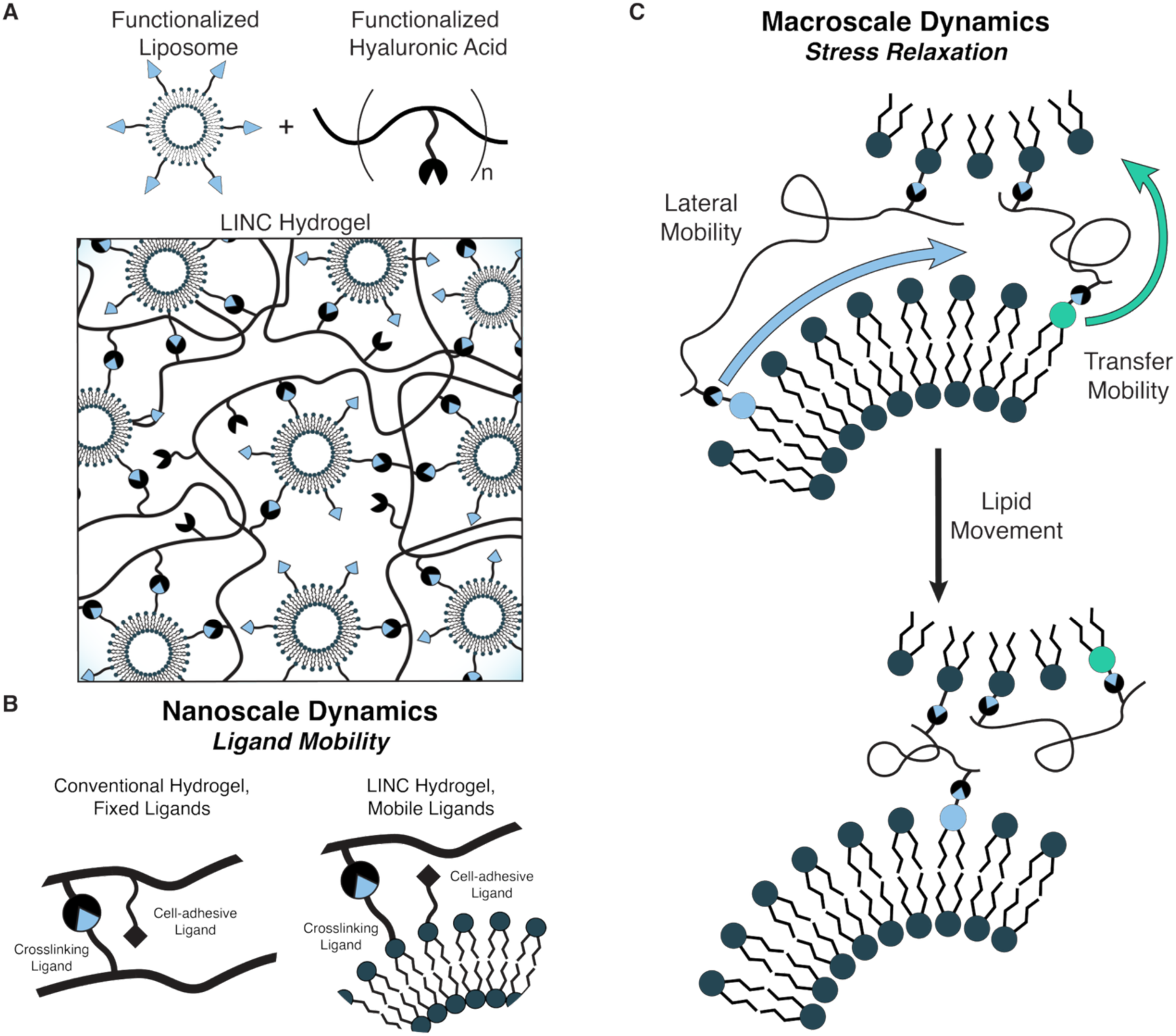
Lipid Network Crosslinked. (**LINC**) **hydrogels.** (**A**) Self-assembled liposomes present covalent crosslinking ligands to form a network of hyaluronic acid (**HA**) biopolymers. This design is termed LINC hydrogel. (**B**) Unlike conventional viscoelastic hydrogels where functional ligands are fixed to the polymer backbone, LINC hydrogels offer control over nanoscale dynamics by tethering ligands to the mobile lipid bilayer. Examples of functional ligands include crosslinking functional groups (circles) or cell-adhesive peptides (black diamonds). (**C**) Due to both the lateral mobility and transfer mobility of the lipid-tethered crosslinking ligands, LINC hydrogels readily relax in response to applied stress, offering control over macroscale network dynamics.

## 2. Results and Discussion

### 2.1 Covalent LINC gels enable control of macroscale stress relaxation

We hypothesized that the dynamic nature of lipid self-assembly would afford stress relaxation properties to covalently crosslinked gels, which typically would be quasi-elastic due to their permanent crosslinking points [6]. To test this, we formulated LINC gels with permanent, static covalent bonds between HA biopolymers and self-assembled liposomes using norbornene and tetrazine functional groups (termed LINC Static, **Figure 2A, Figure S1**). As a comparative control, we synthesized a conventional hydrogel composed only of HA biopolymers using identical permanent, static covalent crosslinks (termed HA Static, **Figure 2A**). The degree of functionalization was controlled to ensure the two formulations had identical HA concentration (1 wt%) and similar stiffness (plateau shear storage modulus, *G’* ∼ 200 Pa; **Figure 2B**).

**Figure 2.**
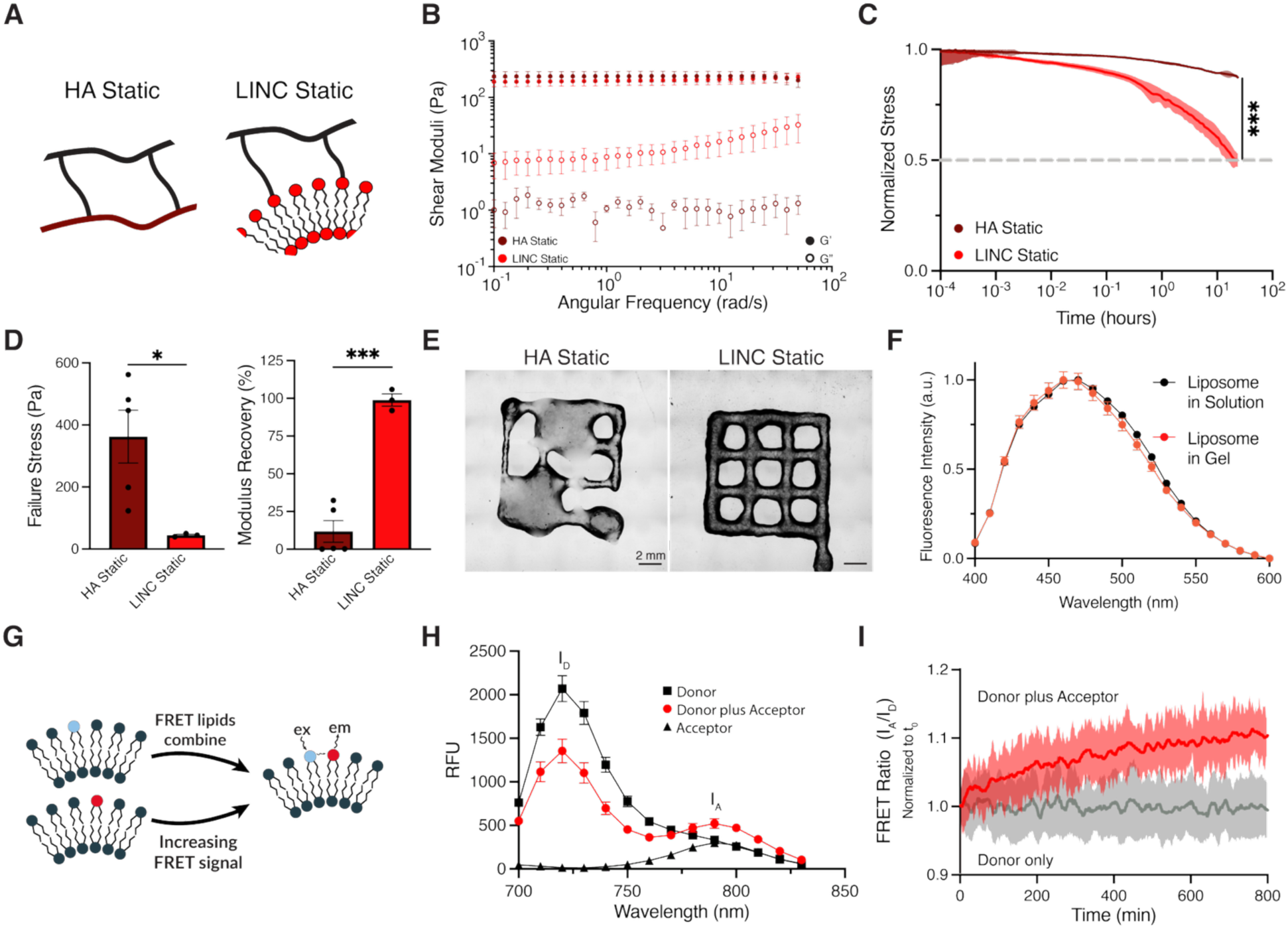
Lipid mobility generates stress relaxation despite irreversible crosslinking in static covalent LINC gels. (**A**) Schematic of HA-only gel and LINC gel with static covalent crosslinks. (**B**) Rheology frequency sweeps of LINC Static and HA Static gels. Shear storage (*G’*) and loss (*G”*) moduli are filled and open symbols, respectively (N=3-5 independent replicates, mean ± standard deviation). (**C**) Normalized stress relaxation of LINC Static and HA Static gels (N=3 independent replicates, mean ± standard deviation, ***p=0.0003 at 24 hours, two-tailed unpaired *t* test). (**D**) Failure stress and plateau shear storage modulus recovery post failure. **(E)** 3D printing of “open window” lattice tests. The lipid mobility significantly increases the yielding behavior and the ability to recover mechanical properties after yielding, thus making the gels printable. **(F)** LAURDAN emission spectra for liposomes in solution (black) and crosslinked into LINC Static gels (red) showing the lipid membrane maintains fluidity when crosslinked. N=3, data are mean ± standard deviation. **(G)** Schematic of FRET experiment showing lipid transfer across liposomes. (**H**) Fluorescence spectra of donor only, acceptor only, and donor plus acceptor LINC Static gels. **(I)** Increasing FRET ratio over time confirms lipid transfer between liposomes, compared to the donor only control (N=3 independent replicates, mean ± standard deviation).

Consistent with literature on chemical hydrogels [6,15], HA Static gels demonstrated limited stress relaxation, appearing quasi-elastic over 24 hours. In contrast, LINC Static hydrogels reached 22% relaxation after only 1 hour and 50% relaxation after 24 hours (**Figure 2C**). Furthermore, LINC Static gels exhibited lower failure stress and greater recovery post failure, making them injectable and suitable as inks for extrusion-based 3D printing, while HA Static gels were unable to be printed (**Figure 2D-E, Figure S5)**. LINC Static gels presented higher loss moduli than HA Static gels at every measured frequency, indicating a larger viscous contribution (**Figure 2B**). This suggests that lipid mobility within the self-assembled liposomes enables viscous dissipation in LINC gels despite the irreversibility of the static covalent crosslinking. Thus, the network harnesses the molecular motion of individual lipids to enable network-level stress relaxation.

### 2.2 Lipids maintain nanoscale mobility in LINC gels

We reasoned that the self-assembled lipids could contribute to viscous dissipation and macroscale stress relaxation through two different modes: (i) lateral mobility within the membrane of a single liposome and (ii) transfer mobility from one liposome to another (**Figure 1C**). Both motions would enable the covalently-bound HA chains to reorient and dissipate applied stress. To confirm both modes of lipid movement occur in LINC gels, we first compared the overall membrane fluidity of liposomes in solution to those crosslinked into a gel. The fluorescent dyes 6-acetyl-2- dimethylaminonaphthalene (ACDAN) and 6-dodecanoyl-2-dimethylaminonaphthalene (LAURDAN) are sensitive to local water, which correlates with lipid membrane fluidity [31,32].

The fluorescence of ACDAN, which positions itself closer to the water interface, and for LAURDAN, which partitions further into the lipid bilayer due to a longer acyl tail, were similar for liposomes in solution and those in LINC gels, suggesting the overall membrane fluidity is retained after network crosslinking (**Figure 2F, Figure S6**).

After confirming that lipids are still mobile after being crosslinked into a hydrogel, we next examined potential lipid movement between liposomes using Förster Resonance Energy Transfer (FRET) (**Figure 2G-I**). Liposomes labeled only with donor dye (Cy5.5) or only with acceptor dye (Cy7) were mixed with functionalized HA to create a LINC Static gel (**Figure 2G**). Upon excitation of the donor, the FRET signal increased over time as individual lipids transferred into the same liposomes, moving the donors and acceptors close together (Förster radius ∼5-7 nm [33,34], **Figure 2H, I**). As expected, no changes were observed in donor-only and acceptor-only controls (**Figure 2I, Figure S7**). Overall, these results confirm that lipids maintain their membrane mobility and can transfer across liposomes while crosslinked into larger polymeric networks, validating the hierarchical, biomimetic design strategy.

### 2.3 Dynamic covalent LINC gels exhibit tunable macroscale viscoelasticity

To demonstrate versatility in these hierarchical materials, we next modified the crosslinking ligands to enable DCC crosslinking as an alternative to static covalent crosslinking. DCC crosslinking has emerged as a strategy to design chemical hydrogels with tunable control of macroscale stress relaxation rates; however, these systems typically have relatively slow stress relaxation half-lives compared to hydrogels with non-covalent crosslinks [12,35]. We reasoned that combining DCC with the lipid mobility of LINC gels would result in rapidly stress-relaxing gels due to combined contributions from both the chemical crosslinks and lipid movement. To test this idea, we modified HA biopolymers with a benzaldehyde motif as previously described [15]. Liposomes were decorated with hydrazine groups that react with benzaldehyde to form a dynamic covalent hydrazone bond with appreciable on- and off-rates at physiological conditions [12,36,37].

We compared the viscoelasticity of these hydrazone-crosslinked gels (termed LINC Dynamic) to HA-only hydrogels (termed HA Dynamic) prepared with the same DCC crosslinks (**Figure 3A, Figure S2-4**). After mixing benzaldehyde-modified HA with either hydrazine-modified HA or hydrazine-modified liposomes, hydrogels rapidly formed within 5-10 seconds (**Figure S8**). As before, the degree of modification was selected to achieve similar gel stiffness (*G’* ∼ 200 Pa) with an equivalent concentration of HA (1 wt%) (**Figure 3B**). Both HA Dynamic and LINC Dynamic formulations displayed shear thinning, self-healing, and strain stiffening behavior, which are common in DCC crosslinked gels (**Figure S9**) [38]. Importantly, upon application of a step strain, the HA Dynamic gels exhibited a stress relaxation half-life (t_1/2_) of 14.2 hours, while t_1/2_ for LINC Dynamic gels was an order of magnitude faster at 1.2 hours (**Figure 3C**). In conventional DCC hydrogels, polymer chains move when the dynamic bonds are off to dissipate stress, making stress relaxation dependent on crosslinking reaction kinetics [35]. Here, by combining lipid mobility with reversible covalent crosslinking, we achieved a ∼12-fold increase in stress relaxation rate despite identical crosslinking chemistry.

**Figure 3.**
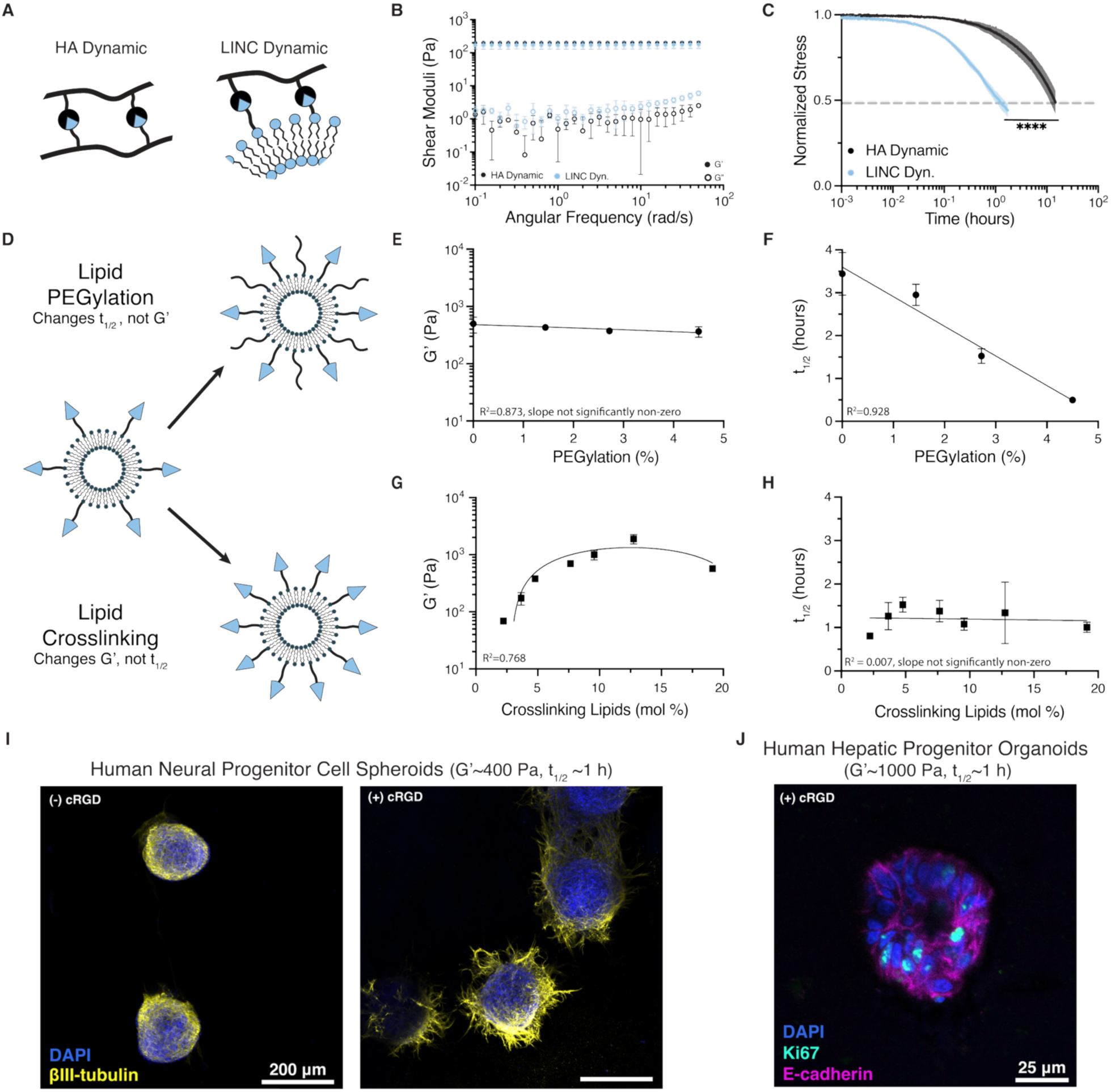
Combining lipid mobility with dynamic covalent crosslinking results in tunable viscoelasticity. (**A**) Schematic of HA-only and LINC gels with dynamic covalent crosslinks. (**B**) Rheology frequency sweeps of LINC Dynamic and HA Dynamic gels. Shear storage (*G’*) and loss (*G”*) moduli are filled and open symbols, respectively (N=3-5 independent replicates, mean ± standard deviation). (**C**) Normalized stress relaxation of LINC Dynamic and HA Dynamic gels (N=3 independent replicates, mean ± standard deviation. ****p<0.0001, two-tailed unpaired *t* test). (**D**) Schematic showing control of LINC gel viscoelasticity through modifying liposome PEGylation (top) or crosslinking density (bottom). (**E-F**) The resulting storage modulus (E) and t_1/2_ (F) of LINC Dynamic gels with increasing PEGylation. (N=3-5 independent replicates, fits are linear regressions). (**G-H**) The resulting storage modulus (G) and t_1/2_ (H) of LINC Dynamic gels with increasing crosslinker functionalization (N=2 independent replicates for 19 mol %, N=3-5 for all others, mean ± standard deviation, fits are quadratic (G) and linear regression (H)). All comparisons in (E, H) not statistically significant (p≥0.32 (E); p≥0.27 (H)) using ordinary one-way ANOVA with Tukey’s multiple comparison test. (**I**) Human neural progenitor cell spheroids cultured for 7 days in LINC Dynamic gels (G’ ∼400 Pa, t_1/2_ ∼1 h) without (left) and with (right) lipid-anchored cRGD. Scale bar is 200 µm. (**J**) Human hepatic progenitor organoids cultured in LINC Dynamic gels (G’ ∼1000 Pa, t_1/2_ ∼1 h) with lipid-anchored cRGD for 7 days. Scale bar is 25 µm.

We next explored strategies to tune the viscoelastic properties of LINC hydrogels through liposome design (**Figure 3D**). We hypothesized that displaying polyethylene glycol (PEG) from the liposome surface would modulate lipid mobility, and hence stress relaxation rate, without impacting the degree of crosslinking. Separately, we hypothesized that altering the concentration of crosslinking lipids would tune the resulting gel stiffness without impacting lipid mobility. When combined, these two strategies allow us to precisely control the viscoelastic properties of LINC gels.

To test the first hypothesis, we varied the fraction of PEGylated lipids, which is expected to increase lipid membrane fluidity [39], while holding the amount of hydrazine functionalization approximately constant. The resulting LINC gels all had similar stiffness (*G’* ∼ 400 Pa) (**Figure 3E**). As expected, increasing PEGylation allowed for faster stress relaxation rates, with an inverse linear relationship between t_1/2_ and the percentage of PEGylated lipids (R^2^ = 0.928) (**Figure 3F**). Compared to LINC Dynamic gels without PEGylation, adding PEG to 4.5% of lipids decreased t_1/2_ by about one order of magnitude to 30 minutes, ∼20-fold faster than the HA Dynamic gels (**Figure 3F**) and approaching reported values of soft tissues [3,40].

To test the second hypothesis, we fixed the PEGylation to hydrazine ratio and varied the concentration of hydrazines available for crosslinking. A maximum stiffness (G’) of ∼1,900 Pa was achieved for LINC Dynamic gels with 12.75 mol% crosslinking lipids (**Figure 3G**). Further increases in crosslinking lipid resulted in lower stiffness, presumably due to defects such as HA wrapping around individual liposomes instead of bridging across particles. Importantly, the t_1/2_ stayed largely constant for all gel formulations (**Figure 3H**), despite the storage modulus changing by an order of magnitude. Thus, by leveraging the nanoscale design of the liposome lipid bilayers, we were able to independently tune the hydrogel stiffness and the stress relaxation rate, which is difficult to achieve with conventional chemical gels without modifying crosslinking chemistry.

Tunable control of hydrogel mechanical properties is particularly helpful when designing *in vitro* models of human tissues [3,15,36], so we next explored if these materials could serve as scaffolds to support the growth of 3D cultures. Human pluripotent stem cell-derived 3D tissue models, such as organoids and spheroids, are increasingly used to recapitulate native tissues and elucidate mechanisms of development, disease, and therapeutic drug effects [41,42]. To demonstrate the modularity and cytocompatibility of the LINC Dynamic platform, we formulated gels to have stiffnesses known to be supportive of human neural progenitor cell (hNPC) spheroids or hepatic progenitor organoids, *G’* ∼ 400 or 1,000 Pa, respectively [43,44]. To engage with integrin cell-surface receptors, we designed the liposomes to present lipid-anchored, cell-adhesive ligands (**Figure 1B**). Specifically, we presented cyclic RGD (cRGD) peptides, a common cell-adhesive ligand that mimics an integrin epitope found in the native extracellular matrix [45]. To confirm bioactivity of cRGD-presenting liposomes, induced pluripotent stem cell-derived hNPCs were encapsulated as spheroids in LINC Dynamic gels with and without the cRGD ligand (1.5 and 0 mM cRGD, respectively). Without cRGD, hNPC spheroids failed to extend βIII-tubulin-positive neurites, while spheroids in LINC Dynamic gels with lipid-anchored cRGD exhibited robust neurite extension (**Figure 3I**). For the culture of human hepatic progenitor organoids, we designed a stiffer gel with 1.5 mM cRGD and laminin (1 mg/ml), as these matrix properties are reported to support hepatic organoids [44]. Hepatic progenitors derived from human induced pluripotent stem cells were encapsulated as single cells within LINC Dynamic gels and cultured 7 days to form organoids (**Figure 3J**). The hepatic progenitor organoids showed high viability, expression of epithelial markers (E-cadherin), and maintenance of proliferation (Ki67+) (**Figure 3J, Figure S10**). Taken together, these data demonstrate that LINC Dynamic gels are cytocompatible and modular to support the culture of different human cell types.

### 2.4 Programming macroscale and nanoscale dynamics in LINC gels

Next, we sought to program the nanoscale dynamics in LINC gels by leveraging our control over the physicochemical properties of the self-assembled liposomes. Thus far, all LINC gel formulations consisted of liposomes with exclusively unsaturated lipids. Lipid tail saturation is a well-known factor influencing the characteristics of lipid bilayers [4]. We hypothesized that saturated lipids would have slower membrane mobility due to their denser packing, offering a means to control both macroscale viscoelasticity and nanoscale ligand mobility. To demonstrate, we designed liposomes with exclusively saturated lipids or mixed liposomes with both saturated and unsaturated lipids (**Figure 4A**).

**Figure 4.**
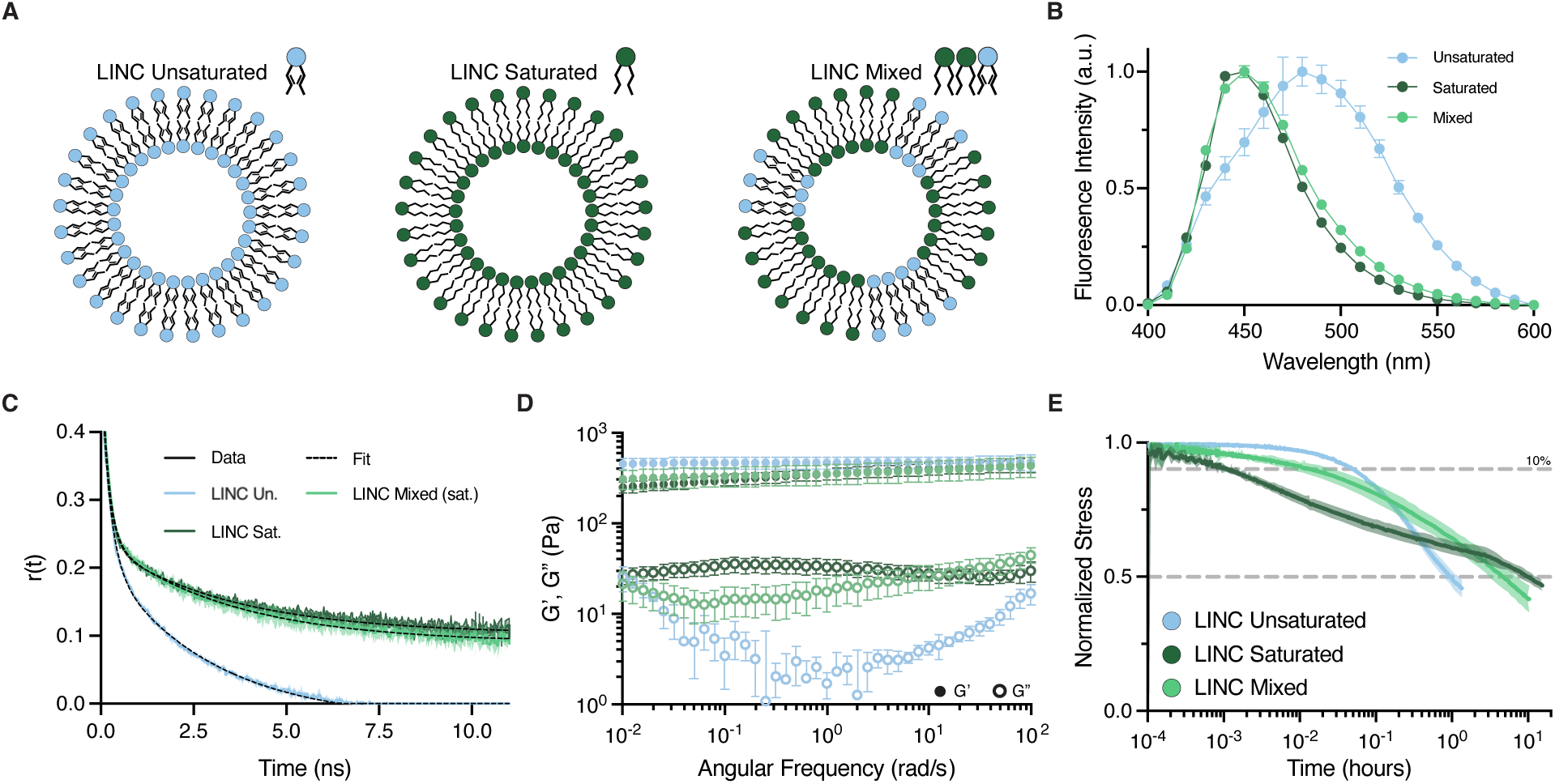
Lipid saturation can program nanoscale and macroscale LINC gel dynamics. (**A**) Schematic of liposomes composed of unsaturated, saturated, or a 1:2 mix of unsaturated and saturated lipids. (**B**) LAURDAN fluorescence spectra of LINC Unsaturated, Saturated, and Mixed gels (N=3 independent replicates, mean ± standard deviation). (**C**) Time resolved fluorescence anisotropy decay in liposomes composed of unsaturated, saturated, or 1:2 unsaturated:saturated mix (N=3 independent replicates, mean ± standard deviation. Fits are two-term exponential decays). **(D)** Oscillatory rheology frequency sweep of LINC Unsaturated, Saturated, and Mixed gels. N=6-7 independent hydrogels. Data are mean ± standard deviation. (**E**) Stress relaxation of LINC Unsaturated, Saturated, and Mixed gels (N=6-7 independent replicates, mean ± standard deviation).

At physiological conditions, saturated liposomes exist in a solid-like gel phase with restricted bilayer fluidity [4]. As expected, the fully saturated liposomes displayed lower mobility compared to fully unsaturated liposomes, as confirmed by LAURDAN/ACDAN membrane fluidity measurements and differential scanning calorimetry (DSC) (**Figure 4B, Figure S11**). To characterize individual lipid-level dynamics, we used time-correlated single photon counting (TCSPC) fluorescence anisotropy decay to study the relative movement of fluorescently labeled lipids within liposomes in different gel conditions. This revealed that the fluorescence anisotropy decay in the LINC Saturated liposomes was notably slower than in LINC Unsaturated, with a slower initial decay and a higher residual anisotropy (**Figure 4C, Figure S12**). Taken together, these data confirm that lipid movement is more restricted within the LINC Saturated liposomes compared to the LINC Unsaturated liposomes.

We hypothesized that this slower lipid mobility would also result in slower stress relaxation at the larger network-level length scale when crosslinking ligands were tethered to saturated lipids. Consistent with this idea, changing tail saturation led to a notable change in loss modulus frequency response, while storage moduli were relatively unchanged (**Figure 4D**). Interestingly, while t_1/2_ was significantly greater for LINC Saturated gels compared to stiffness-matched LINC Unsaturated gels (**Figure 4E**, 10.3 hours *vs* ∼1 hour, respectively), the stress relaxation profile was not uniformly slowed. Instead, at shorter times (less than ∼5 min) the stress relaxation occurred more rapidly in the LINC Saturated gels, with tissue-like kinetics [1,40]. For example, the time required to relax 10% of the applied stress was more than an order of magnitude faster for LINC Saturated compared to LINC Unsaturated, (**Figure 4E, Figure S13**, 5 *vs* 183 s, respectively). We rationalized that the initial rapid relaxation may be due to the reduced random lipid motion caused by inherent thermal energy, which may enable greater directional bias of early lipid movement upon application of a directional stress. However, at longer times, long-range lipid motion is restricted by the limited membrane fluidity, resulting in the expected slower t_1/2_. Thus, this hierarchical design of nanoscale self-assembly within a macroscale polymer network enables the programming of complex viscous behavior such as a bimodal stress relaxation profile.

We reasoned that combining unsaturated and saturated lipids within individual liposomes would enable us to control the macroscale stress relaxation profile and nanoscale ligand dynamics separately by selecting which lipid phase the respective ligands are presented from. For mixed liposomes (33% Unsaturated, 67% Saturated lipids), self-assembly into discrete lipid domains with high and low mobility within single liposomes is expected to occur. DSC confirmed that mixed liposomes exhibit two phases, indicative of intraparticle lipid phase separation (**Figure S11**). We designed the liposomes to present crosslinking ligands from the more mobile unsaturated lipids, while the less mobile saturated lipids presented the cell-adhesive ligands. This enables the modular design of macroscopic stress relaxation profile and nanoscale ligand mobility within one gel, where we use one population of lipids for network crosslinking and a second population for cell adhesion. As an estimate of membrane fluidity in the saturated domains of LINC Mixed gels, the fluorescence of LAURDAN (which has a long, saturated acyl tail) was comparable to LINC Saturated gels (**Figure 4B**). Similarly, TCSPC fluorescence anisotropy showed similar anisotropy decays of labeled saturated lipids for LINC Mixed and LINC Saturated gels, which were notably slower than LINC Unsaturated (**Figure 4C, Figure S12**). These data demonstrate that ligands displayed from saturated lipids have similar local nanoscale mobility in both LINC Mixed and LINC Saturated gels.

We then assessed the macroscopic network stress relaxation of LINC Mixed gels. As expected, since the unsaturated lipids were used as crosslinking points, the stress relaxation rate at both short and long time-scales was intermediate to the LINC Unsaturated and Saturated gels (10% and 50% relaxation at 65 s and 5.3 hours, respectively) (**Figure 4E, Figure S13**). Thus, we can modulate the intraparticle lipid composition to control the macroscopic network stress relaxation dynamics while maintaining nanoscale mobility. The result is a LINC Mixed gel with similar nanoscale ligand dynamics to the LINC Saturated gels, but significantly differing macroscale viscoelasticity.

### 2.5 Human neural progenitor cells respond to both macroscale and nanoscale network dynamics

We next used this system to investigate the consequences of both macroscale stress relaxation and nanoscale cell-adhesive ligand mobility on hNPC morphology (**Figure 5**). hNPCs were encapsulated as single cells (and not as pre-formed spheroids as in Figure 3) to increase their cellular surface contact area with the engineered matrix and to decrease potential confounding signaling from cell-cell interactions inherent to cellular spheroids. The bioactive cRGD ligand was conjugated to either the HA biopolymer, a common method of ligand presentation, or saturated lipids in either the LINC Saturated or Mixed gels. Bioactive ligand presentation was found to have no impact on macroscale viscoelasticity for any LINC gel formulation (**Figure S14**). In all formulations, liposome size, liposome concentration, HA concentration, HA benzaldehyde functionalization, gel stiffness, and total cRGD concentration were kept constant (**Table S1, S2**). After encapsulation within the LINC gels, hNPCs were cultured for 7 days and then analyzed for development of βIII-tubulin-positive neuritic projections and overall morphology (**Figure 5**). To evaluate the role of nanoscale cell-interactive ligand mobility within gels of identical macroscale viscoelasticity, hNPCs were cultured in LINC Saturated gels with the bioactive cRGD fixed either to the HA biopolymer or to individual lipids within the saturated liposomes (**Figure 5A-C**). When the cRGD ligand was fixed to the HA biopolymer, hNPCs displayed modest βIII-tubulin expression with neurites mostly restricted to large cellular spheroids with limited extension into the surrounding matrix (**Figure 5B, C**). These spontaneously forming cellular aggregates are commonly referred to as neurospheres, and this observation is consistent with previous reports that NPCs cultured in gels with longer t_1/2_ display increased neurosphere formation [46]. Interestingly, when the presentation of the cRGD ligand was moved to the liposomes in LINC Saturated gels, we observed a dramatic increase in βIII-tubulin expression (**Figure 5B**). In these gels, the hNPCs primarily remained as single cells (**Figure 5C**) with long, extended neurites averaging ∼200 microns (**Figure 5A, F**). These results are consistent with two-dimensional lipid bilayer studies of nanoscale ligand mobility, where cell-interactive ligand presentation via high viscosity membranes enhances cell receptor engagement [47,48]. Thus, for the stress relaxation profile of the LINC Saturated gels, the local nanoscale mobility of bioactive ligands within the three-dimensional network significantly impacts cell phenotype.

**Figure 5.**
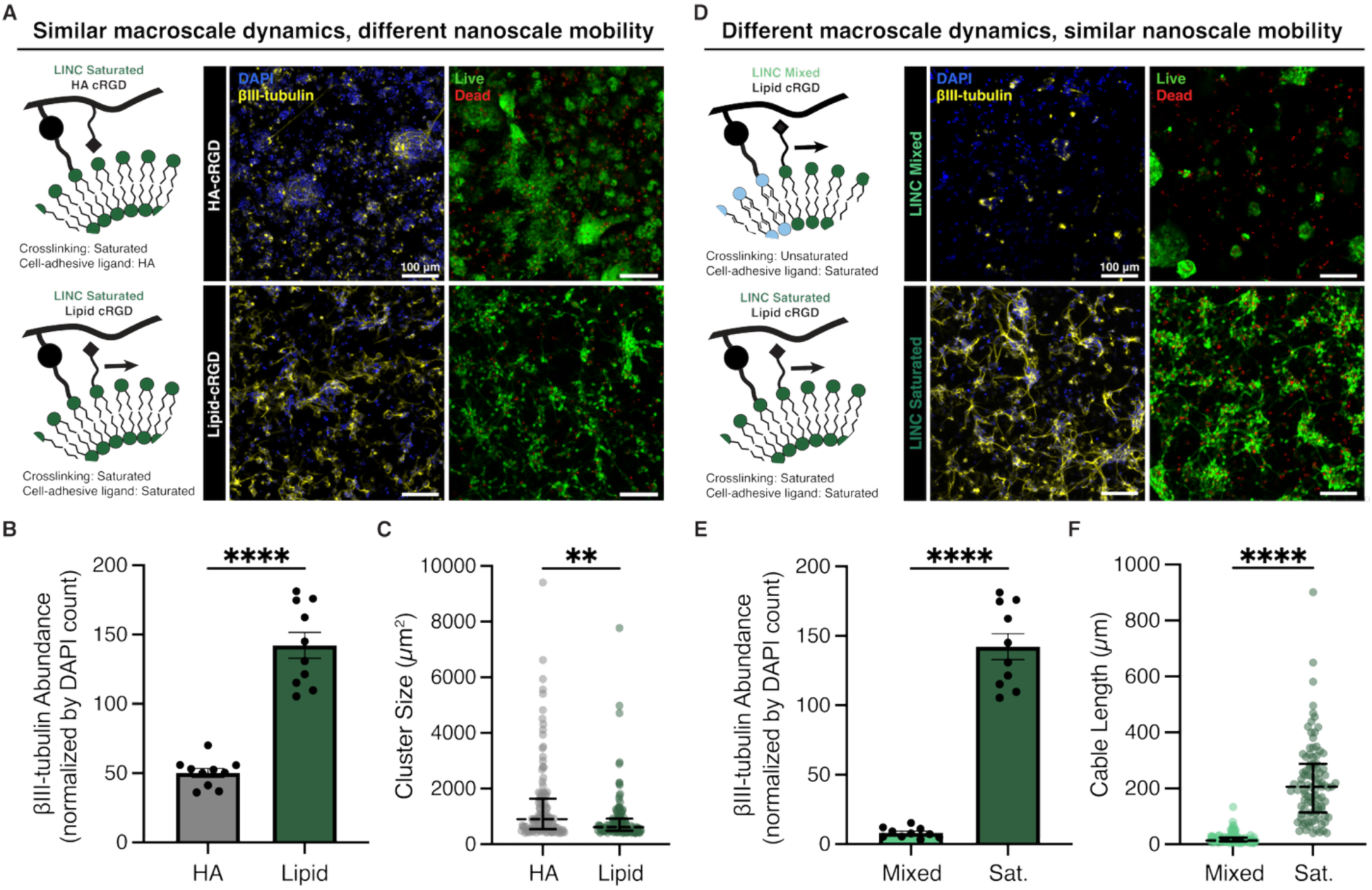
LINC gel design reveals that hNPCs respond differentially to macroscale and nanoscale dynamics. (**A**) Representative fluorescence images of hNPCs encapsulated as single cells in LINC Saturated gels with either lipid- or HA-anchored cRGD after 7 days of culture with labeled nuclei (DAPI, blue) and neuritic extensions (βIII-tubulin, yellow). Scale bar is 100 µm. (**B**) Quantification of ýIII-tubulin area normalized by DAPI count in LINC Saturated HA- or lipid-presented cRGD. (N=3 replicate hydrogels, n=5 fields of view, mean ± standard deviation. ****p<0.0001, 2-way ANOVA with Tukey’s multiple comparisons test.) (**C**) Quantification of hNPC cluster size. (N=3 replicate hydrogels, n=3-4 fields of view. Data points are individual cell clusters. Bars are medians with interquartile range. ****p<0.0001, Kruskal-Wallis with Dunn’s multiple comparison test.) (**D**) Representative fluorescence images of hNPCs encapsulated as single cells in LINC Saturated and Mixed gels with lipid-anchored cRGD after 7 days of culture with labeled nuclei (DAPI, blue) and neuritic extensions (ýIII-tubulin, yellow). Scale bar is 100 µm. (**E**) Quantification of ýIII-tubulin area normalized by DAPI count in LINC Saturated and Mixed with lipid-presented cRGD. (N=3 replicate hydrogels, n=5 fields of view, mean ± standard deviation. ****p<0.0001, 2-way ANOVA with Tukey’s multiple comparisons test.) (**F**) Quantification of hNPC neurite cable length in lipid-anchored-cRGD LINC gels. (N=3 replicate hydrogels, n=3-4 fields of view. Data points represent single cells with traceable neurites. Bars are medians with interquartile range. ****p<0.0001, Kruskal-Wallis with Dunn’s multiple comparison test).

Finally, to explore the effect of macroscale viscoelasticity in gels of similar nanoscale dynamics on hNPC phenotype, we cultured hNPCs in LINC Saturated and Mixed gels with the cRGD ligand presented on saturated lipids in both conditions (**Figure 5D-F**). While both cultures remained primarily as single cells, LINC Mixed gels did not support the appreciable expression of βIII-tubulin (**Figure 5E**) nor the extension of neuritic projections (**Figure 5F**). This indicates that when the cRGD is presented by the mobile lipid phase, the initial rate of network stress relaxation is at least as impactful as the oft reported t_1/2_. Taken together, these results highlight the context dependent effects of both macroscopic stress relaxation dynamics and nanoscale ligand mobility and emphasize the critical need for hydrogels with programmable control of network dynamics across multiple length scales

## 3. Conclusion

Here we introduce the use of self-assembled lipids as dynamic structural elements in engineered hydrogels. Crosslinking through individual, mobile lipids within liposomes provides a new approach to control stress relaxation by leveraging bond mobility. With equivalent crosslinking chemistries, lipid network crosslinked hydrogels show 20-fold faster stress relaxation than polymer-only gels. We demonstrate that liposome surface modification, lipid saturation, and mixed phase lipid membranes provide control over the short and long timescales of stress relaxation. Lipid-bound ligand presentation is tuned independently of viscoelasticity through these same design principles. Finally, we demonstrate the importance of this unique hierarchical control by showing human neural progenitor cells exhibit significantly distinct responses to macroscale stress relaxation depending on the nanoscale dynamics of ligand mobility. These results motivate future biophysical studies of the molecular pathways and mechanisms that enable cells to respond to different time-scales and length-scales of hierarchical network dynamics, and provide a platform to do so.

Looking ahead, the large body of literature on liposome formulations suggests that this class of materials will have a large design space to explore structure-function relationships to achieve a broad range of multi-length-scale dynamics. The LINC hydrogel strategy is compatible with any chemically modifiable biopolymer, allowing for future application-specific designs of self-assembled lipids with covalent polymer crosslinks. While here the liposomes were used solely as dynamic structural elements, they could simultaneously be exploited for their more traditional roles as drug delivery vehicles or extracellular vesicle signals ^[21–24,26,49]^. When incorporated into hydrogels, self-assembled lipids are lubricating [50], enhance water retention in flexible sensors [51], and serve as tunable reservoirs for drug release [52]. While we did not explore these properties here, we hypothesize LINC gels will show similar characteristics. Overall, this work establishes LINC hydrogels as a versatile system using liposomes as intentional structural elements with programmable control over both macroscale and nanoscale dynamics.

## 4. Methods

### Lipids

| Lipid | Vendor |
| --- | --- |
| DOPC | Broadpharm BP-25710 |
| DOPE-mal | Broadpharm BP-28874 |
| 18:1 DBCO PE | Avanti Polar Lipids 870129 |
| DSPC | Broadpharm BP-25623 |
| DSPE-mal | Broadpharm BP-26159 |
| 16:0 DBCO PE | Avanti Polar Lipids 870128 |
| 16:0 NBD PE | Avanti Polar Lipids 810144 |
| 18:1 NBD PE | Avanti Polar Lipids 810145 |
| 18:1 Cyanine 5.5 PE | Avanti Polar Lipids 810336 |
| 18:1 Cyanine 7.0 PE | Avanti Polar Lipids 810337 |

### Statistical Analysis

All statistical analysis was performed using GraphPad Prism.

### HA Benzaldehyde Synthesis

HA Benzaldehyde with a 20% modification was synthesized as previously described [15]. First, HA is modified with an alkyne group via a carbodiimide reaction to form the intermediate HA-Alkyne. 100 kDa HA (sodium hyaluronate, LifeCore Biomedical) is fully dissolved at 1 wt% in 2-(N-morpholino)ethanesulfonic acid (MES) buffer [0.2 M MES hydrate (Sigma) and 0.15 M NaCl in Milli-Q water (pH 4.5)]. Then, 3 equivalence propargylamine (Sigma) per HA dimer unit was added and the solution was adjusted to pH 6.0 with NaOH. N-hydroxysuccinimide (3 eq., Thermo Fisher Scientific) was then added as a powder. Immediately after, 3 eq. 1-ethyl-3-(3-dimethylaminopropyl)carbodiimide hydrochloride (EDC) (Sigma) was added as a powder and the reaction was stirred overnight at room temperature (RT). The reaction was dialyzed against Milli-Q water for 3 days, filtered, lyophilized, and stored at -20 °C.

The HA-Alkyne was then reacted with an azidobenzaldehyde molecule via a copper-catalyzed azide-alkyne click chemistry reaction to form HA Benzaldehyde (HA-BZA). The lyophilized HA-alkyne was dissolved in 10x phosphate-buffered saline [PBS; 81 mM sodium phosphate dibasic, 19 mM sodium phosphate monobasic, and 60 mM sodium chloride in Milli-Q water (pH 7.4)] with β-cyclodextrin (1 mg/ml; Sigma-Aldrich) at 1 wt%. The dissolved HA-alkyne was degassed under nitrogen for 30 minutes. Sodium ascorbate (0.18 eq., Sigma-Aldrich) and copper (II) sulfate pentahydrate (0.0096 eq., Sigma-Aldrich) were dissolved in Milli-Q water and degassed under nitrogen for 30 minutes prior to adding them to the HA-alkyne solution. 4-Azidobenzaldehyde (3eq., Santa Cruz Biotechnology) was dissolved in anhydrous DMSO and added to the HA-Alkyne, sodium ascorbate, and copper (II) sulfate pentahydrate solution. The reaction was stirred for 24 hours at RT. Then, an equal volume of EDTA was added for 1 hour to chelate the copper (50 mM, pH 7.0, Thermo Fisher Scientific). The reaction was dialyized against Milli-Q water for 3 days, filtered, lyophilized, and stored at -20 °C.

For cRGD functionalization, HA-BZA was further reacted with Cyclo(-RGDfk) (MedChemExpress) to form HA-BZA cRGD. The same EDC/NHS reaction as used to prepare the HA-alkyne was used with 3 eq of c(-RGDfk) per HA dimer unit. ^1^H NMR spectra were recorded using nuclear magnetic resonance (NMR) spectroscopy (Varian Inova, 600 MHz) using deuterated D_2_O as a solvent.

### HA Hydrazine Synthesis

First, N-(3-azidopropyl)-2-hydrazineylacetamide was synthesized as previously described [53]. Briefly, tri-Boc-hydrazinoacetic acid (1 eq.), azidopropyl amine (1.3 eq.), 1-ethyl-3-(3-dimethylaminopropyl)carbodiimide (1.3 eq.) (EDC), and 4-dimethylaminopyridine (0.2 eq.) were dissolved in methylene chloride (DCM) and stirred overnight. Excess solvent was removed using a rotary evaporator under reduced pressure, and the remaining product was purified via silica gel chromatography. The Boc-protected product was then dissolved in a 1:1 mix of DCM:trifluoroacetic acid (TFA) for 4 hours to deprotect and precipitated in ether to form an oily yellow solid. Next, HA modification occurred in the same manner as for HA Benzaldehyde, as described above, except the synthesized N-(3-azidopropyl)-2-hydrazineylacetamide was used in place of 4-Azidobenzaldehyde. 1 eq. of propargylamine, EDC, NHS, and -(3-azidopropyl)-2-hydrazineylacetamide were used per HA dimer unit. ^1^H NMR spectra were recorded using nuclear magnetic resonance (NMR) spectroscopy (Varian Inova, 600 MHz) using deuterated D_2_O as a solvent.

### HA Tetrazine and HA Norbornene Synthesis

HA was modified with tetrazine or norbornene as previously described [15]. Unmodified HA (100 kDa, LifeCore Biomedical) was fully dissolved in 0.1 M MES buffer (pH 7) at 1 wt%. Then, 1-hydrozybenzotriazole hydrate (2 eq. per HA dimer unit; Sigma-Aldrich) was added and dissolved for 15 minutes. Separately, tetrazine amine (2 eq. per HA dimer unit; Conju-Probe) or norbornene amine (2 eq. per HA dimer unit; TCI America) were dissolved in acetonitrile (MeCN) mixed with deionized water (5:1). EDC (2 eq) was added as a powder and allowed to dissolve. The mixture was added dropwise to the dissolved HA over 30 min and reacted overnight at room temperature. The reaction mixture was dialyzed for 1 day against a 10% MeCN solution, followed by 3 days against Milli-Q water. The resulting product was then sterile-filtered, lyophilized, and stored at -20°C. ^1^H NMR spectra was recorded using NMR spectroscopy (Varian Inova, 600 MHz) using deuterated D_2_O as a solvent.

### SH-PEG-Hydrazine Synthesis

SH-PEG2k-NH_2_ was dissolved in dimethyl formamide (DMF) at 75 mg/mL. Separately, tri-Boc-hydrazinoacetic acid (Sigma, 3 eq.) in equal volume DMF, 4-methylmorpholine (NMM, Sigma, 5 eq.), and HATU (Sigma, 3 eq.) were added sequentially, with 15 minutes in between each. The tri-Boc-hydrazinoacetic acid containing solution was let stir for 1 hour. Then, the SH-PEG-NH_2_ (1 eq.) was added dropwise to the tri-Boc-hydrazinoacetic acid mixture at roughly 1 mL / min and left to react for 20-24 hours at RT. The reaction was precipitated in ice-cold diethyl ether 3x and dried overnight under nitrogen gas and subsequently dried under vacuum for 3 days. The collected pellet was dissolved in a solution of 50:50 DCM:TFA for 1 hour to remove the Boc group. The product was again precipitated 3x in diethyl ether and dried. Of note, deprotection times longer than 1 hour led to inconsistent results. Though we did not confirm it, we suspect this is due to TFA-mediated cleavage of the amide bond that has been reported in similar hydrazine-containing molecules [54]. ^1^H NMR spectra were recorded using nuclear magnetic resonance (NMR) spectroscopy (Varian Inova, 600 MHz) using deuterated D_2_O as a solvent.

### SH-PEG-Norbornene Synthesis

First, SH-PEG2k-NH_2_ was dissolved at approximately 200 mg/mL in anhydrous DCM. Then, 4-Dimethylaminopyridine (1 eq. to NH_2_ DMAP, Sigma-Aldrich) and exo-5-Norbornenecarboxylic acid (6 eq., Sigma-Aldrich) were added as powders. After dissolving, EDC (6 eq.) was added as a powder. The mixture was degassed under nitrogen for 20 minutes and the reaction proceeded for 24 hours at 4 °C in the dark. The reaction was precipitated in ice-cold diethyl ether 3x and dried overnight under nitrogen gas and subsequently dried under vacuum for 3 days. Finally, the dried product was dissolved in Milli-Q water, filtered through a 0.22 µm filter, and lyophilized prior to use. ^1^H NMR spectra were recorded using nuclear magnetic resonance (NMR) spectroscopy (Varian Inova, 600 MHz) using deuterated D_2_O as a solvent.

### Liposome fabrication

Liposomes were prepared using a standard thin film hydration and sonication method. Briefly, lipids in chloroform were mixed at the desired ratios (**Table S1**) and dried under nitrogen gas then left under vacuum overnight to remove excess chloroform. The resulting lipid films were hydrated in PBS for 10 minutes with slight. This solution was then sonicated until ∼100 nm diameter particles were formed. For liposome formulations containing saturated lipids, this process was done at 70 °C, above the highest reported T_m_ of the lipids being used. As the used unsaturated lipids are above their T_m_ at room temperature, no heating was required.

Once 100 nm particles were achieved, 1.25 eq. of the crosslinking linker (SH-PEG2k-Hydrazine, SH-mPEG, or SH-PEG2k-Norbornene) was added for 1 hour at 37 °C for functionalization. Lastly, the functionalized particles were purified using 100 kDa amicon spin filter units (Millipore Sigma). Purification consisted of 3 PBS washes followed by concentrating to a final concentration of 0.17 M, or approximately 133 mg lipids/mL PBS, assuming no loss in the fabrication process. Liposome size was verified via dynamic light scattering (DLS).

### Hydrogel formation

For HA-HA gels, all HA conditions were dissolved at 1 wt% in 1x PBS. Once dissolved, the complementary HAs were mixed at a 1:1 volume ratio and hydrogels spontaneously formed.

For LINC Gels, all HA conditions were dissolved at 2 wt% HA in 1x PBS. Once purified, the liposomes were mixed at a 1:1 volume ratio with the functionalized HA for a final concentration of 1 wt% HA. Liposomes were at a final concentration of approximately 67 mg/mL in all hydrogel conditions. Crosslinking occurs spontaneously upon mixing. Liposomes were used within 2 days of fabrication.

### Rheological characterization

Rheological measurements were carried out using an ARG2 rheometer (TA Instruments) with a 20 mm, 1° cone-on-plate geometry. Mineral oil was used to seal the hydrogels and prevent dehydration during measurement. Hydrogels were cast at 23 °C then the temperature was ramped to 37 °C at 2 °C/min. Time sweeps were conducted at 1% oscillatory strain and 1 Hz frequency and allowed to continue until a plateau in moduli was observed. Frequency sweeps were performed from 0.01 to 100 rad/s under 1% oscillatory strain. The reported G’ and G” values were taken at 6.28 rad/s (1 Hz) from the frequency sweeps. To further characterize the viscoelasticity, stress relaxation measurements were performed under a constant strain of 5%. For stress relaxation in Figure 2C , data is smoothed with a 2nd order Gaussian filter over 15 data points due to noise at long times.

Stress sweeps were conducted to determine the failure stress and post-failure recovery of the gels. They were performed with steady state sensing from 1 Pa to failure, as defined by the measured strain exceeding 1000%, indicative of the geometry rotating freely post gel failure as all gel conditions fail at strains under 1000%. Reported failure stress values were defined as the last point prior to the above failure definition, also accompanied by an abrupt drop in viscosity. Modulus recovery post failure was defined by G’ 5 minutes after the termination of the failure stress measurement using a time sweep at 1% oscillatory strain and 1 Hz frequency. The shear-thinning and self-healing measurements were conducted by performing flow peaks holding for 30 s at alternative low (0.1 s^-1^) and high (10 s^-1^) shear rates. All measurements were performed at 37 °C except for the initial time sweeps.

### 3D printing

Suitability of the LINC Static inks for extrusion-based 3D (bio-)printing was investigated using a MakerGear M2 3D printer equipped with a multi-material extrusion system [55] and a straight 27-gauge dosing needle (inner diameter 210 µm). Gels were cast and allowed to gel within the syringe body for 2 hours prior to use. Square-shaped lattice structures (9x9 mm^2^) consisting of four perpendicular layers and a strand distance 3 mm were printed with constant extrusion rate and a printing speed of 400 mm/min. The prints were imaged with a Leica THUNDER fluorescence microscope.

### Förster Resonance Energy Transfer (FRET)

Liposomes were prepared as described above. For each condition, 0.1 mol% of the appropriate fluorescent lipids in chloroform were added during the thin-film drying process. Cy5.5 was used as the donor while Cy7 was used as the acceptor. Donor liposomes were prepared with 0.1 mol% Cy5.5, acceptor liposomes with 0.1 mol% Cy7, and unlabeled liposomes with no fluorophores.

LINC hydrogels were cast as described above into a black, flat bottom 96 well plate (Costar). Immediately post casting, the samples were excited at 675 nm (donor excitation) and read at both 720 nm (donor emission) and 790 nm (acceptor emission) every minute. Conditions consisted of 1) donor and acceptor lipids on different liposomes (Donor plus Acceptor), 2) donor liposomes mixed with unlabeled liposomes as a control (Donor), and 3) acceptor liposomes mixed with unlabeled liposomes as a control (Acceptor). The donor-only and acceptor-only conditions contained the same amount of the respective fluorescent lipids as the D+A condition. The reported FRET Ratio is the ratio of the acceptor fluorescence intensity (790 nm) to the donor fluorescence intensity (720 nm) at each timepoint, normalized to the first time point (t_0_). Spectra were collected from 700-850 nm at the end of each measurement (ex 675 nm). PBS filled wells were used as blanks. Data is smoothed with a 2nd order Gaussian filter over 5 data points.

### Differential Scanning Calorimetry (DSC)

Liposomes were prepared as described. Liposome solutions in PBS were placed in hermetically sealed TA instruments T zero pans. An empty pan was used as a reference. Samples were cooled from room temperature to -40 °C, held at - 40 °C for 5 minutes, and heated to -5 °C and held for 5 minutes. This was repeated twice. Samples were then heated to 23°C, held for 5 minutes, heated to 80°C and held for 5 minutes, cooled to 23 °C and held. This was repeated twice. Temperatures were ramped at a rate of 2 °C min^-1^.

### ACDAN/LAURDAN

Liposomes were prepared as stated with the addition of 0.05 mol% of either ACDAN (Santa Cruz Biotech) or LAURDAN (Lumiprobe) dissolved in chloroform during the thin film drying process. 40 µL gels or liposomes at the same concentration were cast in a 96 well plate. The samples were excited at 360 nm and readings from 400-600 nm were taken 2 hours post casting, once the gels were approximately fully crosslinked. PBS filled wells were used as blanks.

### Time correlated single photon counting fluorescence anisotropy

Measurements were conducted using a Horiba Fluorolog-3 Spectrofluorometer with an automated polarizer L-format. Liposomes were prepared as stated with the addition of 0.2 mol% 18:1 NBD PE (Avanti) or 16:0 NBD PE (Avanti). Measurements were taken at an excitation of 460 nm and emission of 535 nm. G factor correction was conducted with free fluorophore and held constant. IRF correction was performed using 100 nm PS beads (Polysciences). Liposomes were diluted 1:1200 to an approximate concentration of 0.11 mg/mL in PBS. Liposome concentration, slit size, and laser power were all held constant. Fitting was performed using the Horiba DAS-6 analysis software.

### Human neural progenitor cell (hNPC) culture

The stemness and differentiation capacity of all hiPSCs was previously validated [56]. Approval for this study was obtained from the Stanford Institutional Review Board, and informed consent was obtained from all donors. hiPSCs were cultured with mTeSR-1 Plus (STEMCELL Technologies) media in monolayer on human embryonic stem cell (hESC)–qualified Matrigel (0.1 mg/mL; Sigma-Aldrich) to maintain their pluripotent state.

hNPCs were differentiated from hiPSCs and cultured as previously described [15]. Briefly, a commercially available dual SMAD inhibition kit (STEMCELL Technologies) was used to differentiate hiPSCs into hNPCs. hiPSCs were expanded to confluency on hESC-qualified Matrigel before being exposed to STEMdiff Neural Induction Medium for 10 days with daily media changes. After, cells were dissociated with cell dissociation solution (Sigma-Aldrich), passaged onto poly-d-lysine (50 μg/mL; Sigma-Aldrich) and laminin (10 μg/mL; Roche) coated plates. At this point, hNPCs were maintained in N3 culture medium consisting of DMEM/F12 (Thermo Fisher Scientific), neurobasal (Thermo Fisher Scientific), N-2 supplement (1%, Thermo Fisher Scientific), B-27 supplement with vitamin A (2%, Thermo Fisher Scientific), GlutaMax (1%, Thermo Fisher Scientific), and Minimal Essential Medium Nonessential Amino Acids (1%, Thermo Fisher Scientific).

### hNPC encapsulation

hNPCs were either encapsulated as single cells or formed into spheroids prior to encapsulation. For spheroid encapsulation, hNPCs were dissociated and about 1.5 × 10^6^ single cells were seeded per well of an AggreWell 800 plate (STEMCELL Technologies) in N3 media. After 48 hours, hNPC spheroids of approximately 5,000 cells were then collected, pelleted by centrifugation, and resuspended in a 2 wt% HA-BZA solution. Gels were cast and allowed to gel for 30 min. 750 µL of N3 media was added to each well containing a 10 µL hydrogel. Medium was changed daily.

For single cell encapsulation, hNPCs were dissociated, filtered through a 70 µm cell strainer, pelleted by centrifugation, and counted. They were then resuspended in a 2 wt% HA-BZA solution at a cell density of 6x10^4^ cells/µL, for a final gel concentration of 3 × 10^4^ cells/µL. Gels were cast and allowed to gel for 30 min. 750 µL of N3 media was added to each well containing a 10 µL hydrogel. Medium was changed daily.

### Manual neurite tracing

bIII-tubulin-positive neurites were manually traced and quantified with the SNT toolbox to assess their length and branching complexity [57]. Neurites with a total length less than 5 μm were not traced; otherwise, to the greatest degree possible, all neurites longer than 5 μm were traced in 3D space.

### Cell viability

Cell-containing hydrogels were washed with DPBS and incubated with 2 µM calcein acetoxymethyl and 4 µM ethidium homodimer for 20 min at 37 °C. The gels were washed with DPBS and imaged with a confocal microscope (Leica SPE).

### Hepatic progenitor organoid generation and passaging in Matrigel

Hepatic organoids were derived from iPSCs using previously reported methods [44]. Human iPSC-derived hepatic progenitor cells were cultured in 50 µL Matrigel domes at 200 cells/µL seeding density within a 24-well plate to generate hepatic organoids. Organoids were passaged every 7 days. To passage organoids, Matrigel domes were incubated for 30 min with 1 mL ice-cold, 5 mM ethylenediamine tetra-acetic acid (EDTA) in PBS to dissociate the gels. The cells were then collected and centrifuged for 5 min at 500 x g and treated with 1 mL TrypLE (Thermo Fisher Scientific) for 6 min at 37 °C. Gentle mixing by pipette aspiration every 3 min assisted the organoid dissociation into single cells. The TrypLE was then quenched with 400 µL 40% FBS in PBS, and centrifuged for 5 min at 500 x g. The cell pellet was resuspended in hepatic growth medium for cell counting. The desired number of cells were centrifuged for 5 min at 500 x g and resuspended in fresh, ice-cold Matrigel solution at 200 cells/µL. The Matrigel domes were incubated for 10 min of gelation at 37 °C, then 700 µL of pre-warmed organoid growth medium was added to each well. Y-27632 (10 µM Cayman Chemical) was added to the medium for the first 2 days. Media was replaced every 2–3 days. To make complete hepatic growth media, RPMI media was supplemented with 1x B27 (Thermo), 250 nM LDN-193189 (Cayman Chemical), 3 µM CHIR99021 (Cayman Chemical), 10 µM A83-01 (Cayman Chemical), 100 ng/mL EGF (Cayman Chemical), 10 ng/mL FGF10 (Cayman Chemical), 20 ng/mL HGF (Cayman Chemical).

### Hepatic progenitor organoid culture in LINC gels

HA and liposome stock solutions were prepared as described above. 6 mg/mL laminin-111 (R&D Systems) was premixed with the liposome solution to have a final concentration of 1 mg/mL in LINC gels. Hepatic organoids were released from Matrigel and dissociated into single cells as described above. The desired number of cells were centrifuged for 5 min at 500 x g and resuspended in the solution of liposomes and laminin. The gels were cast as previously stated, with a cell seeding density of 2 k/µL. Gels were cast and allowed to gel for 30 min. Following gelation, LINC hydrogels were submerged in organoid passage media (growth media with Y-27632) for 2 days, then changed to growth media with changes every 2-3 days for a week.

### Immunocytochemistry (ICC)

Importantly, ICC was performed using Tween-20 as a permeabilization agent instead of Triton X-100 as Triton X-100 disrupts the liposome structure, leading to immediate gel dissociation even post fixation.

Cell-containing hydrogels were washed with DPBS, fixed in a 4% paraformaldehyde solution for 20 min at 37 °C, followed by 3 washes with DPBS. Then, samples were permeabilized for 1 hour at room temperature with 0.1% (v/v) Tween-20 in PBS (PBST) and subsequently blocked with PBS supplemented with 5% (w/v) bovine serum albumin (BSA, Roche), 5% (v/v) goat serum (Gibco), and 0.1% (v/v) Tween-20 for 3 hours at room temperature with gentle rocking. Primary antibodies were prepared in PBS with 2.5% (w/v) BSA, 2.5% (v/v) goat serum, and 0.1% (v/v) Tween-20 (antibody dilution solution). Mouse anti-β3-tubulin was used at a 1:400 dilution (Cell Signaling). Samples were incubated with primary antibodies overnight at 4 °C and then washed with PBS 3 times for 30 min each. Then, secondary antibodies were diluted in antibody dilution solution, added to the samples, and incubated overnight at 4 °C prior to 3 additional PBS washes. Goat anti-mouse Alexa Fluor 488 (Invitrogen) was used at a 1:500 dilution, and diamidino-2-phenylindole (DAPI, 1 µg/mL, Cell Signaling) was included in the secondary antibody solution for nuclear staining. The stained hydrogels were inverted and imaged directly with a confocal microscope (Leica SPE).

Hepatic organoids in gels were fixed with 750 µL of 4% paraformaldehyde (PFA) in DPBS for 15 min. Fixation solution was then removed and three 10 min washes of DPBS were performed. The gels were incubated in 30% sucrose solution at 4 °C overnight. Gels then embedded into a 1 mL 1:1 ratio mixture of 30% sucrose and Tissue-Tek O.C.T Compound (Sakura Finetek USA, Torrance, CA) into Tissue-Tek Cryomold molds (Sakura Finetek USA, Torrance, CA). After for 24 h incubation at RT, samples were moved onto dry ice for rapid freezing (∼ 10 min). The samples were then snap frozen on dry ice and cryo-sectioned into 40 µm sections using a Leica Cryostat instrument.

Sectioned samples were melted at 50 °C for 5-10 min and subsequently washed with DPBS to remove excess O.C.T. The gel sections were circled with lipid-repellent marker pen for further staining. Samples were permeabilized for 1 h with 0.5% v/v Tween 20 in DPBS, then blocked for 3 h in DPBS with 5% v/v goat serum, 0.5% v/v Tween 20 and 5 wt% BSA. Staining was conducted as above with mouse anti-Ki67 was added at 1:200 (Cell Signaling) and rabbit anti E-Cadherin was added to 1:400 (Cell Signaling) primary antibodies. Samples were mounted to No.1 glass over slides with ProLong Gold Antifade Reagent for 24-48 hours at RT in dark. Stained samples were imaged using a confocal microscope (Leica SPE) and analyzed using ImageJ (NIH, v.2.1.0/1.53c).

## Acknowledgments

We thank Dr. Carla Huerta-López and Dr. Fotis Christakopoulos for their helpful feedback and discussions. Part of this work was performed at the Stanford Nano Shared Facilities, supported by the NSF (ECCS-2026822).

## Data availability

All data will be available online in a Stanford University repository upon publication.

## Author Contributions

**Conceptualization:** NJB, SCH **Methodology:** NJB, MSH, NPN, YL, DK, RSN **Investigation:** NJB, MSH, NPN, JAB, JW, YL, RO, DK, RSN **Visualization:** NJB **Funding acquisition:** SCH **Supervision:** SCH **Writing:** NJB, SCH **Manuscript editing:** NJB, MSH, NPN, YL, DK, SCH

## References

1. Chaudhuri, O., Cooper-White, J., Janmey, P.A., Mooney, D.J., and Shenoy, V.B. (2020) Effects of extracellular matrix viscoelasticity on cellular behaviour. Nature, 584 (7822), 535–546.

2. Courbot, O., and Elosegui-Artola, A. (2025) The role of extracellular matrix viscoelasticity in development and disease. *npj Biol*. Phys. Mech., 2 (1), 10.

3. Saraswathibhatla, A., Indana, D., and Chaudhuri, O. (2023) Cell–extracellular matrix mechanotransduction in 3D. Nat Rev Mol Cell Biol, 1–22.

4. Alberts, B., Johnson, A., Lewis, J., Raff, M., Roberts, K., and Walter, P. (2002) The Lipid Bilayer, in Molecular Biology of the Cell. 4th edition, Garland Science.

5. Jahnke, K., and Staufer, O. (2024) Membranes on the move: The functional role of the extracellular vesicle membrane for contact-dependent cellular signalling. Journal of Extracellular Vesicles, 13 (4), e12436.

6. Lou, J., and Mooney, D.J. (2022) Chemical strategies to engineer hydrogels for cell culture. Nat Rev Chem, 6 (10), 726–744.

7. Álvarez, Z., Kolberg-Edelbrock, A.N., Sasselli, I.R., Ortega, J.A., Qiu, R., Syrgiannis, Z., Mirau, P.A., Chen, F., Chin, S.M., Weigand, S., Kiskinis, E., and Stupp, S.I. (2021) Bioactive scaffolds with enhanced supramolecular motion promote recovery from spinal cord injury. Science, 374 (6569), 848–856.

8. 3D bioprinting of dynamic hydrogel bioinks enabled by small molecule modulators | Science Advances.

9. Rijns, L., Baker, M.B., and Dankers, P.Y.W. (2024) Using Chemistry To Recreate the Complexity of the Extracellular Matrix: Guidelines for Supramolecular Hydrogel–Cell Interactions. J. Am. Chem. Soc., 146 (26), 17539–17558.

10. Li, J., and Mooney, D.J. (2016) Designing hydrogels for controlled drug delivery. Nat Rev Mater, 1 (12), 1–17.

11. Hafeez, S., Aldana, A.A., Duimel, H., Ruiter, F.A.A., Decarli, M.C., Lapointe, V., van Blitterswijk, C., Moroni, L., and Baker, M.B. (2023) Molecular Tuning of a Benzene-1,3,5-Tricarboxamide Supramolecular Fibrous Hydrogel Enables Control over Viscoelasticity and Creates Tunable ECM-Mimetic Hydrogels and Bioinks. Advanced Materials, 35 (24), 2207053.

12. McKinnon, D.D., Domaille, D.W., Cha, J.N., and Anseth, K.S. (2014) Biophysically Defined and Cytocompatible Covalently Adaptable Networks as Viscoelastic 3D Cell Culture Systems. Advanced Materials, 26 (6), 865–872.

13. Sousa, V., Ladeira, B., Garanger, E., Lecommandoux, S., Meijer, E.W., Dankers, P.Y.W., Mano, J.F., and Borges, J. (2025) Merging Natural Biopolymers with Supramolecular Chemistry: Emulating the Native Extracellular Matrix’s Complexity. ACS Nano, 19 (33), 29833–29859.

14. Nelson, B.R., Kirkpatrick, B.E., Miksch, C.E., Davidson, M.D., Skillin, N.P., Hach, G.K., Khang, A., Hummel, S.N., Fairbanks, B.D., Burdick, J.A., Bowman, C.N., and Anseth, K.S. (2024) Photoinduced Dithiolane Crosslinking for Multiresponsive Dynamic Hydrogels. Advanced Materials, 36 (43), 2211209.

15. Roth, J.G., Huang, M.S., Navarro, R.S., Akram, J.T., LeSavage, B.L., and Heilshorn, S.C. (2023) Tunable hydrogel viscoelasticity modulates human neural maturation. Sci Adv, 9 (42), eadh8313.

16. Álvarez, Z., Ortega, J.A., Sato, K., Sasselli, I.R., Kolberg-Edelbrock, A.N., Qiu, R., Marshall, K.A., Nguyen, T.P., Smith, C.S., Quinlan, K.A., Papakis, V., Syrgiannis, Z., Sather, N.A., Musumeci, C., Engel, E., Stupp, S.I., and Kiskinis, E. (2023) Artificial extracellular matrix scaffolds of mobile molecules enhance maturation of human stem cell-derived neurons. Cell Stem Cell, 30 (2), 219–238.e14.

17. Diba, M., Spaans, S., Hendrikse, S.I.S., Bastings, M.M.C., Schotman, M.J.G., van Sprang, J.F., Wu, D.J., Hoeben, F.J.M., Janssen, H.M., and Dankers, P.Y.W. (2021) Engineering the Dynamics of Cell Adhesion Cues in Supramolecular Hydrogels for Facile Control over Cell Encapsulation and Behavior. Advanced Materials, 33 (37), 2008111.

18. Rijns, L., Hagelaars, M.J., van der Tol, J.J.B., Loerakker, S., Bouten, C.V.C., and Dankers, P.Y.W. (2024) The Importance of Effective Ligand Concentration to Direct Epithelial Cell Polarity in Dynamic Hydrogels. Advanced Materials, 36 (43), 2300873.

19. Moghaddam, A.S., Dunne, K., Breyer, W., Wu, Y., and Pashuck, E.T. (2025) Hydrogels with multiple RGD presentations increase cell adhesion and spreading. Acta Biomaterialia, 199, 142–153.

20. Huster, D., Maiti, S., and Herrmann, A. (2024) Phospholipid Membranes as Chemically and Functionally Tunable Materials. Advanced Materials, 36 (23), 2312898.

21. Shi, L., Jing, Y., Lu, H., Zhao, F., An, M., Jin, S., Gao, C., Dai, Y., Zhu, Y., Yang, S., Zhang, S., Ye, X., Cai, X., Wang, Y., and Xin, S. (2025) Extracellular Vesicle Crosslinkers Constructing Hydrogels with Stress-Relaxation and Bioactive Protein Modification. Advanced Materials Interfaces, 12 (10), 2400885.

22. Zhang, Q., Wang, Y., Zhu, Z., Ahmed, W., Zhou, D., and Chen, L. (2025) Therapeutic Potential of Injectable Supramolecular Hydrogels With Neural Stem Cell Exosomes and Hydroxypropyl Methylcellulose for Post-Stroke Neurological Recovery. IJN, 20, 2253–2271.

23. Zhang, X., Lyu, Y., Liu, Y., Yang, R., Liu, B., Li, J., Xu, Z., Zhang, Q., Yang, J., and Liu, W. (2021) Artificial apoptotic cells/VEGF-loaded injectable hydrogel united with immunomodification and revascularization functions to reduce cardiac remodeling after myocardial infarction. Nano Today, 39, 101227.

24. Liang, Y., and Kiick, K.L. (2016) Liposome-Cross-Linked Hybrid Hydrogels for Glutathione-Triggered Delivery of Multiple Cargo Molecules. Biomacromolecules, 17 (2), 601–614.

25. Mouritsen, O.G. (2011) Lipids, curvature, and nano-medicine. European Journal of Lipid Science and Technology, 113 (10), 1174–1187.

26. Correa, S., Grosskopf, A.K., Klich, J.H., Hernandez, H.L., and Appel, E.A. (2022) Injectable liposome-based supramolecular hydrogels for the programmable release of multiple protein drugs. Matter, 5 (6), 1816–1838.

27. Talló, K., Bosch, M., Pons, R., Cocera, M., and López, O. (2019) Preparation and characterization of a supramolecular hydrogel made of phospholipids and oleic acid with a high water content.

28. Dessane, B., Smirani, R., Bouguéon, G., Kauss, T., Ribot, E., Devillard, R., Barthélémy, P., Naveau, A., and Crauste-Manciet, S. (2020) Nucleotide lipid-based hydrogel as a new biomaterial ink for biofabrication. Sci Rep, 10 (1), 2850.

29. Akin-Ige, F., and Amin, S. (2024) Stimuli-Responsive Bio-Based Surfactant-Polymer Gels. Colloids and Surfaces A: Physicochemical and Engineering Aspects, 703, 135149.

30. Stancheva, T.N., Georgiev, M.T., Radulova, G.M., Danov, K.D., and Marinova, K.G. (2022) Rheology of saturated micellar networks: Wormlike micellar solutions vs. bicontinuous micellar phases. Colloids and Surfaces A: Physicochemical and Engineering Aspects, 652, 129927.

31. Mangiarotti, A., Siri, M., Tam, N.W., Zhao, Z., Malacrida, L., and Dimova, R. (2023) Biomolecular condensates modulate membrane lipid packing and hydration. Nat Commun, 14 (1), 6081.

32. Orlikowska-Rzeznik, H., Krok, E., Chattopadhyay, M., Lester, A., and Piatkowski, L. (2023) Laurdan Discerns Lipid Membrane Hydration and Cholesterol Content. J. Phys. Chem. B, 127 (15), 3382–3391.

33. Zhang, S., Metelev, V., Tabatadze, D., Zamecnik, P.C., and Bogdanov, A. (2008) Fluorescence resonance energy transfer in near-infrared fluorescent oligonucleotide probes for detecting protein–DNA interactions. Proceedings of the National Academy of Sciences, 105 (11), 4156–4161.

34. Loura, L.M.S. (2012) Simple Estimation of Förster Resonance Energy Transfer (FRET) Orientation Factor Distribution in Membranes. International Journal of Molecular Sciences, 13 (11), 15252–15270.

35. Rizwan, M., Baker, A.E.G., and Shoichet, M.S. (2021) Designing Hydrogels for 3D Cell Culture Using Dynamic Covalent Crosslinking. Advanced Healthcare Materials, 10 (12), 2100234.

36. LeSavage, B.L., Zhang, D., Huerta-López, C., Gilchrist, A.E., Krajina, B.A., Karlsson, K., Smith, A.R., Karagyozova, K., Klett, K.C., Huang, M.S., Long, C., Kaber, G., Madl, C.M., Bollyky, P.L., Curtis, C., Kuo, C.J., and Heilshorn, S.C. (2024) Engineered matrices reveal stiffness-mediated chemoresistance in patient-derived pancreatic cancer organoids. Nat. Mater., 23 (8), 1138–1149.

37. Gilchrist, A.E., Liu, Y., Klett, K., Liu, Y.-C., Ceva, S., and Heilshorn, S.C. (2023) Transient Competitors to Modulate Dynamic Covalent Cross-Linking of Recombinant Hydrogels. Chem. Mater., 35 (21), 8969–8983.

38. de Paiva Narciso, N., Navarro, R.S., Gilchrist, A.E., Trigo, M.L.M., Aviles Rodriguez, G., and Heilshorn, S.C. (2023) Design Parameters for Injectable Biopolymeric Hydrogels with Dynamic Covalent Chemistry Crosslinks. Advanced Healthcare Materials, 12 (27), 2301265.

39. Tabarin, T., Martin, A., Forster, R.J., and Keyes, T.E. (2012) Poly-ethylene glycol induced super-diffusivity in lipid bilayer membranes. Soft Matter, 8 (33), 8743–8751.

40. Chaudhuri, O., Gu, L., Klumpers, D., Darnell, M., Bencherif, S.A., Weaver, J.C., Huebsch, N., Lee, H., Lippens, E., Duda, G.N., and Mooney, D.J. (2016) Hydrogels with tunable stress relaxation regulate stem cell fate and activity. Nature Mater, 15 (3), 326–334.

41. Hofer, M., and Lutolf, M.P. (2021) Engineering organoids. Nat Rev Mater, 6 (5), 402–420.

42. Fatehullah, A., Tan, S.H., and Barker, N. (2016) Organoids as an in vitro model of human development and disease. Nat Cell Biol, 18 (3), 246–254.

43. Roth, J.G., Brunel, L.G., Huang, M.S., Liu, Y., Cai, B., Sinha, S., Yang, F., Pașca, S.P., Shin, S., and Heilshorn, S.C. (2023) Spatially controlled construction of assembloids using bioprinting. Nat Commun, 14 (1), 4346.

44. Liu, Y., Gilchrist, A.E., Johansson, P.K., Guan, Y., Deras, J.D., Liu, Y.-C., Ceva, S., Huang, M.S., Navarro, R.S., Enejder, A., Peltz, G., and Heilshorn, S.C. (2025) Engineered Hydrogels for Organoid Models of Human Nonalcoholic Fatty Liver Disease. Advanced Science, 12 (22), e17332.

45. Ananthanarayanan, B., Little, L., Schaffer, D.V., Healy, K.E., and Tirrell, M. (2010) Neural stem cell adhesion and proliferation on phospholipid bilayers functionalized with RGD peptides. Biomaterials, 31 (33), 8706–8715.

46. Huang, M.S., LeSavage, B.L., Ghorbani, S., Gilchrist, A.E., Roth, J.G., Huerta-López, C., Mozipo, E.A., Navarro, R.S., and Heilshorn, S.C. (2025) Viscoelastic N-cadherin-like interactions maintain neural progenitor cell stemness within 3D matrices. Nat Commun, 16 (1), 5213.

47. Barcelona-Estaje, E., Oliva, M.A.G., Cunniffe, F., Rodrigo-Navarro, A., Genever, P., Dalby, M.J., Roca-Cusachs, P., Cantini, M., and Salmeron-Sanchez, M. (2024) N-cadherin crosstalk with integrin weakens the molecular clutch in response to surface viscosity. Nat Commun, 15 (1), 8824.

48. Bennett, M., Cantini, M., Reboud, J., Cooper, J.M., Roca-Cusachs, P., and Salmeron-Sanchez, M. (2018) Molecular clutch drives cell response to surface viscosity. Proceedings of the National Academy of Sciences, 115 (6), 1192–1197.

49. Ma, Y.-H., Yang, J., Li, B., Jiang, Y.-W., Lu, X., and Chen, Z. (2016) Biodegradable and injectable polymer–liposome hydrogel: a promising cell carrier. Polym. Chem., 7 (11), 2037–2044.

50. Lin, W., Kluzek, M., Iuster, N., Shimoni, E., Kampf, N., Goldberg, R., and Klein, J. (2020) Cartilage-inspired, lipid-based boundary-lubricated hydrogels. Science, 370 (6514), 335–338.

51. Bai, M., Chen, Y., Zhu, L., Li, Y., Ma, T., Li, Y., Qin, M., Wang, W., Cao, Y., and Xue, B. (2024) Bioinspired adaptive lipid-integrated bilayer coating for enhancing dynamic water retention in hydrogel-based flexible sensors. Nat Commun, 15 (1), 10569.

52. Wu, F., Chen, H., Liu, J., and Pang, Y. (2025) Generating structurally and functionally programmable hydrogels by biological membrane hybridization. Nat Protoc, 1–32.

53. Lou, J., Liu, F., Lindsay, C.D., Chaudhuri, O., Heilshorn, S.C., and Xia, Y. (2018) Dynamic Hyaluronan Hydrogels with Temporally Modulated High Injectability and Stability Using a Biocompatible Catalyst. Adv Mater, 30 (22), e1705215.

54. Dalabehera, N.R., Meher, S., Bhusana Palai, B., and Sharma, N.K. (2020) Instability of Amide Bond with Trifluoroacetic Acid (20%): Synthesis, Conformational Analysis, and Mechanistic Insights into Cleavable Amide Bond Comprising β-Troponylhydrazino Acid. ACS Omega, 5 (40), 26141–26152.

55. Pusch, K., Hinton, T.J., and Feinberg, A.W. (2018) Large volume syringe pump extruder for desktop 3D printers. HardwareX, 3, 49–61.

56. Roth, J.G., Muench, K.L., Asokan, A., Mallett, V.M., Gai, H., Verma, Y., Weber, S., Charlton, C., Fowler, J.L., Loh, K.M., Dolmetsch, R.E., and Palmer, T.D. (2020) 16p11.2 microdeletion imparts transcriptional alterations in human iPSC-derived models of early neural development. eLife, 9, e58178.

57. Arshadi, C., Günther, U., Eddison, M., Harrington, K.I.S., and Ferreira, T.A. (2021) SNT: a unifying toolbox for quantification of neuronal anatomy. Nat Methods, 18 (4), 374–377.

