## Supporting Information for "Lipid Network Crosslinked Hydrogels: Controlling Material Dynamics Across Multiple Length Scales Through Lipid Movement"

**List of Supplemental Information**
**Fig. S1**. $^{1}$H NMR spectra for HA-Static.
**Fig. S2.** $^{1}$H NMR spectra for the HA-BZA modification process.
**Fig. S3.** **(A)** $^{1}$H NMR spectra of HA-Hydrazine with **(B)** structural depiction.
**Fig. S4.** $^{1}$H NMR spectra and structures of **(A)** starting SH-PEG2k-$NH_2$, **(B)** SH-PEG2k-Hyd(Boc), and **(C)** SH-PEG2k-Hyd.
**Fig. S5.** 3D printing replicates.
**Fig S6.** Characterization of membrane fluidity.
**Fig. S7.** FRET control condition.
**Fig. S8.** Representative oscillatory rheology time sweeps.
**Fig. S9.** Rheological characterization of strain stiffening, shear-thinning, and recovery.
**Fig. S10.** Cell viability staining in LINC gels.
**Fig. S11.** Membrane fluidity and DSC of LINC Mixed.
**Fig. S12.** Liposome TCSPC fluorescence anisotropy.
**Fig. S13.** LINC gel stress relaxation.
**Fig. S14.** Viscoelasticity of LINC gels with cRGD.

**Table S1.** Summary of liposome formulations for LINC gels.
**Table S2.** Summary of HA characteristics for selected gel conditions.

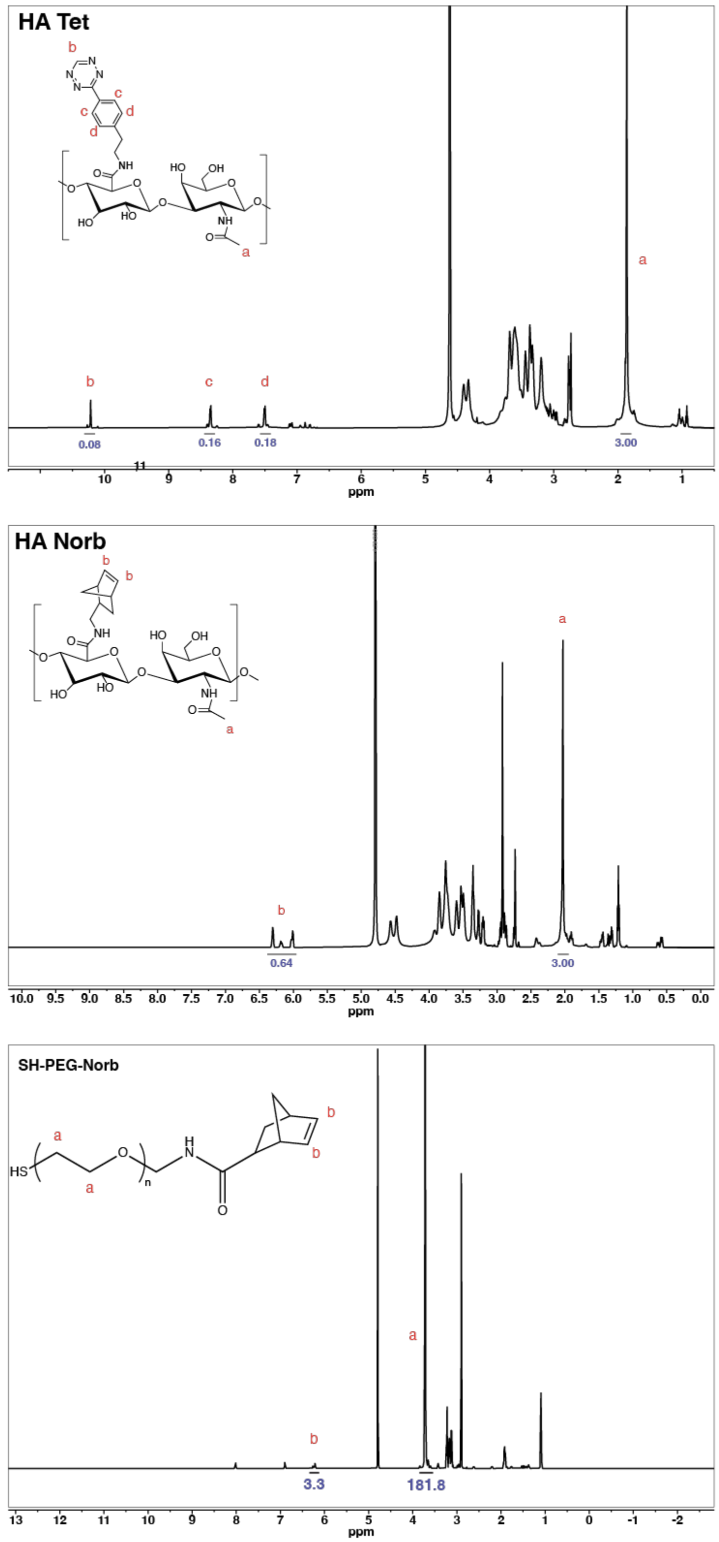


**Fig. S1**. [1]H NMR spectra for HA-Static. Representative spectra for **(A)** tetrazine (8% functionalization) and **(B)** norbornene (32% functionalization) modified hyaluronic acid and **(C)** SH-PEG2k-Norbornene.

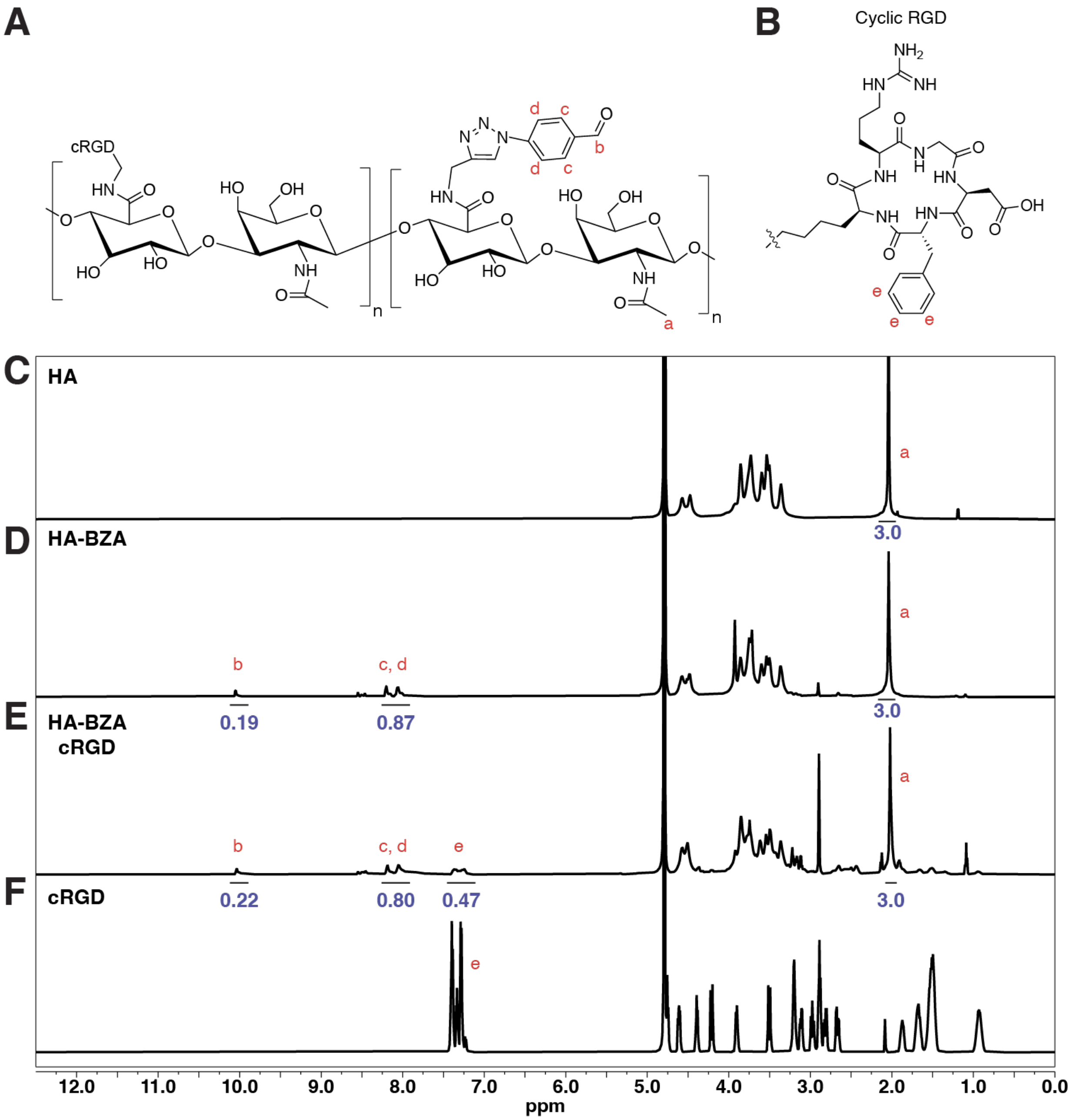


**Fig. S2.** $^{1}$H NMR spectra for the HA-BZA modification process. Structures of **(A)** HA-BZA and **(B)** cyclic RGD peptide. Representative NMR spectra of **(C)** unmodified HA, **(D)** HA-BZA (~20% BZA modification), **(E)** HA-BZA-cRGD (~20% BZA, 15.7% cRGD modification), and **(F)** the starting cRGD ligand.

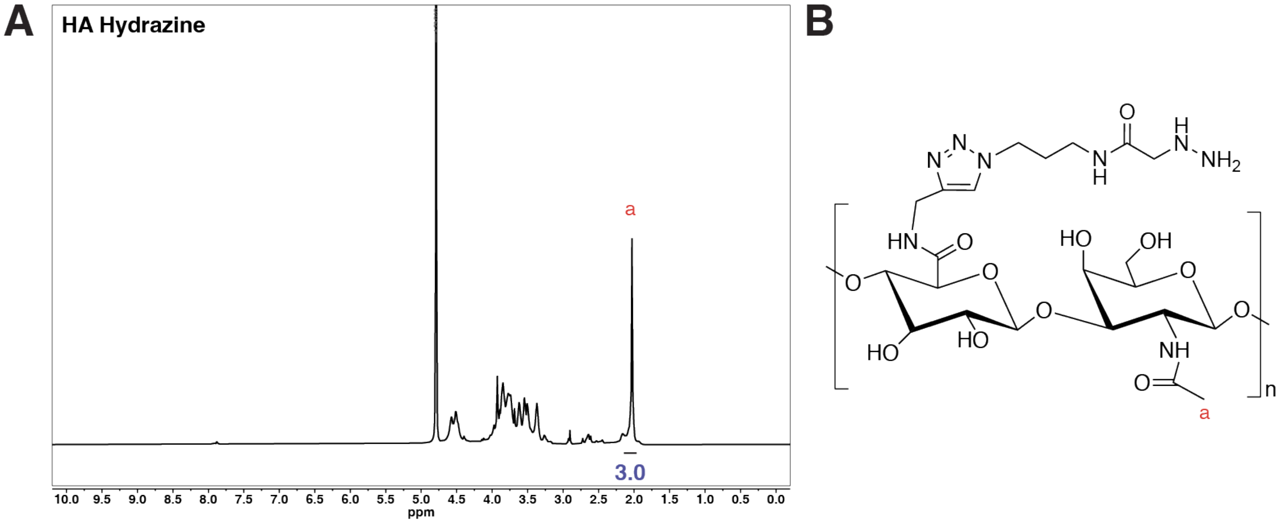


**Fig. S3.** **(A)** $^1$H NMR spectra of HA-Hydrazine with **(B)** structural depiction. Due to peak interference from the HA backbone (see Fig. S2), functionalization was not quantified, and consistency was achieved by evaluating rheological properties of each newly synthesized batch.

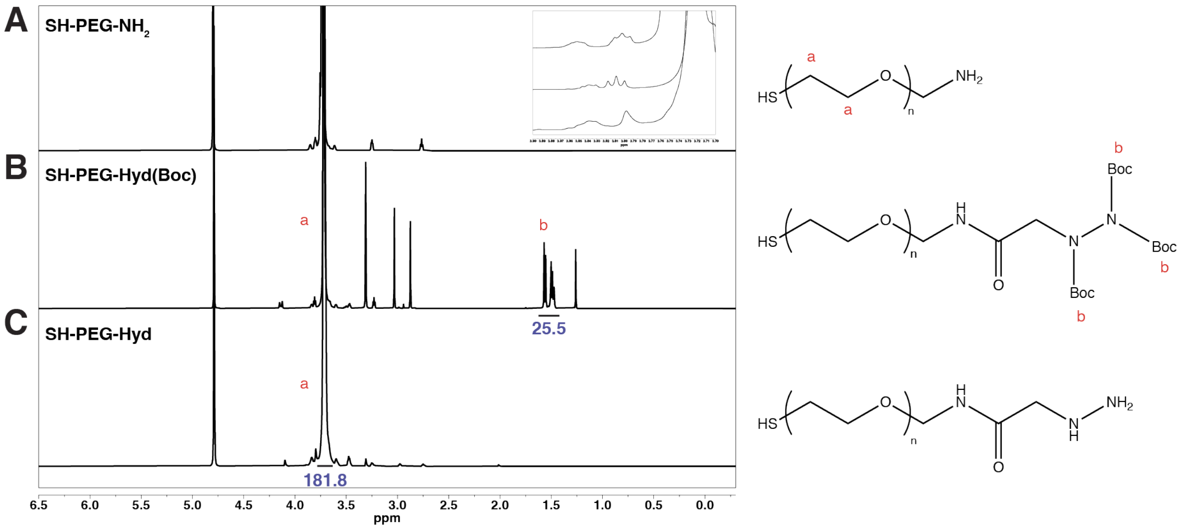


**Fig. S4.** [1]H NMR spectra and structures of **(A)** starting SH-PEG2k-$NH_2$, **(B)** SH-PEG2k-Hyd(Boc), and **(C)** SH-PEG2k-Hyd. Functionalization was calculated to be 94.4% based on Triboc quantification (B).

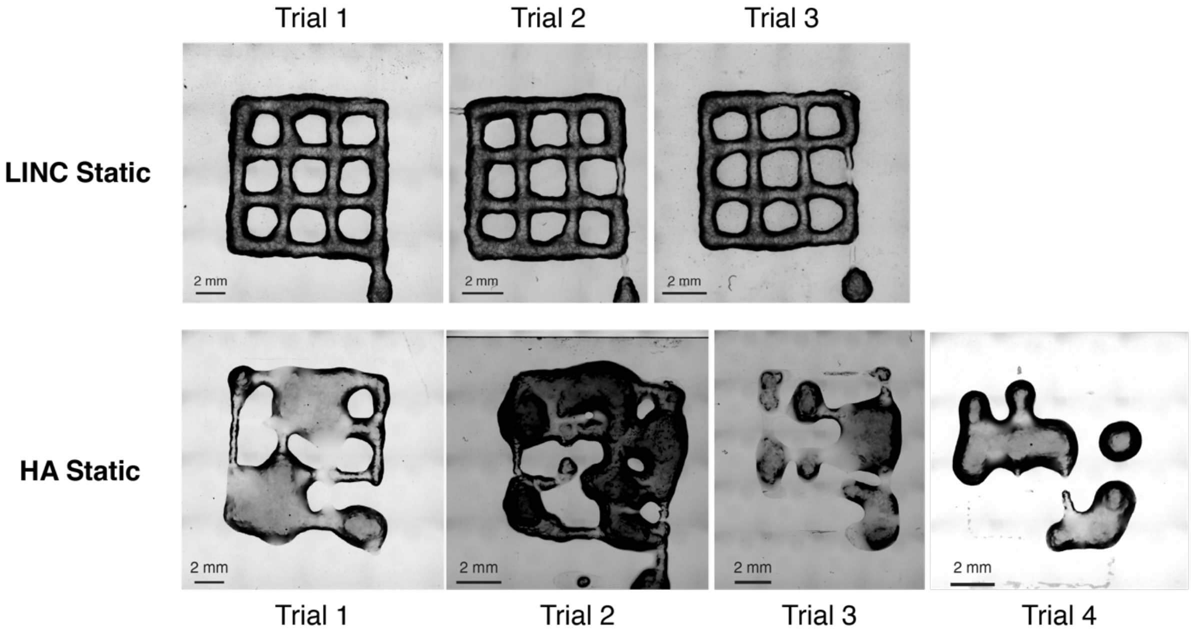


**Fig. S5.** 3D printing replicates. To demonstrate how LINC macroscopic shear-thinning and self-healing network properties enable applications in 3D printing, we tested both formulations with a 28-G printing nozzle. While the HA Static hydrogels were extrudable, their inconsistent flow and frequent, large bursts of material led to them not creating a high-fidelity print. In contrast, the LINC Static hydrogels extruded in a consistent filament with little to no burst, as demonstrated using a common 3D printing "open window" lattice test.

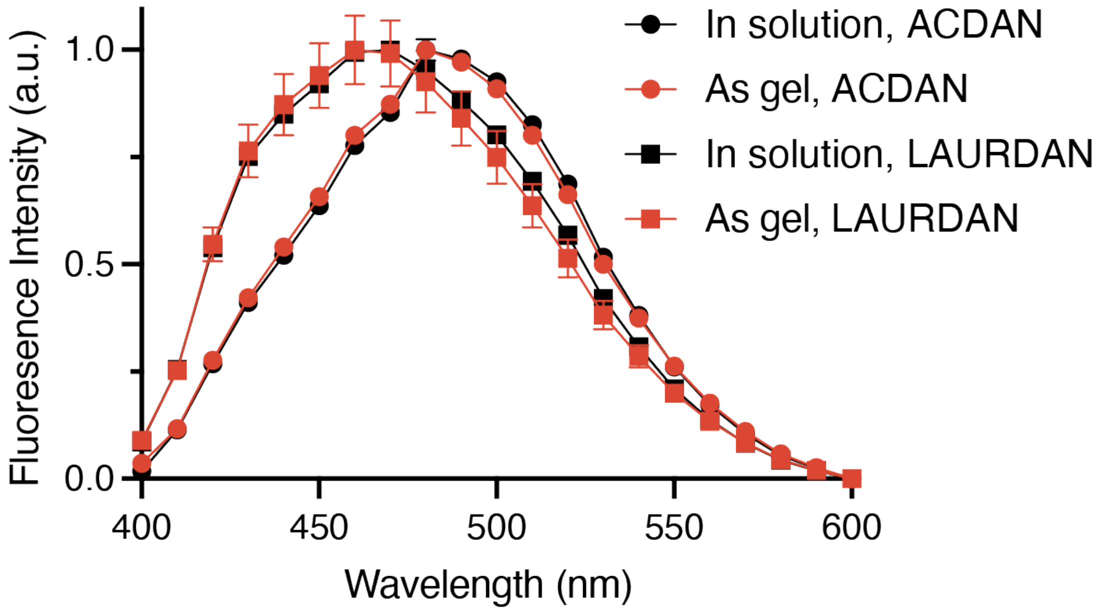


**Fig S6.** Characterization of membrane fluidity. LAURDAN (squares) and ACDAN (circles) emission spectra for liposomes in solution (black) and crosslinked into LINC Static gels (red). N=3, data are mean ± standard deviation.

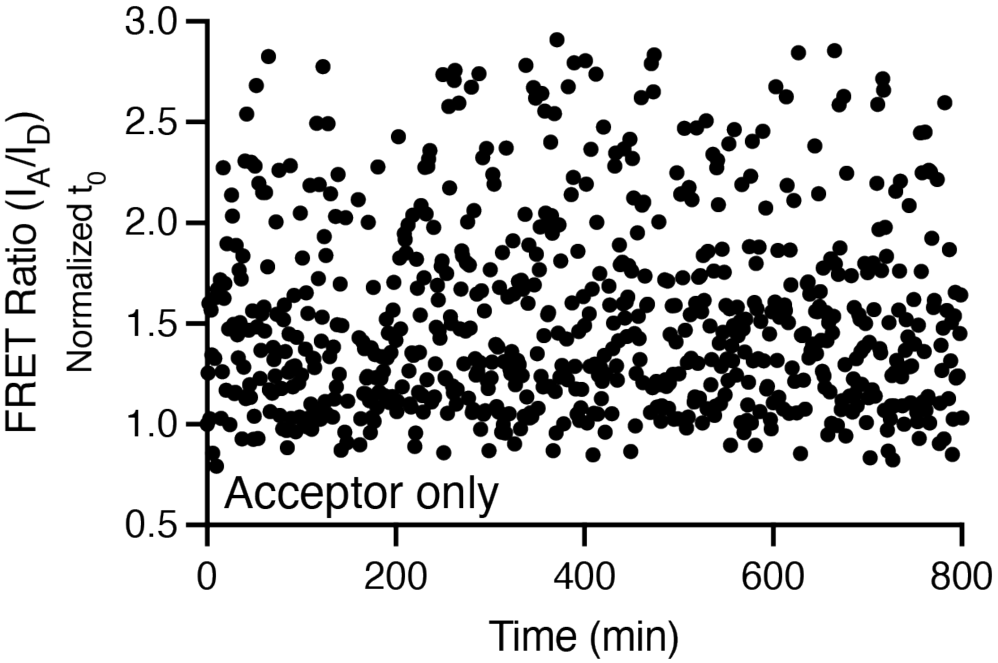


**Fig. S7.** FRET control condition. The FRET Ratio of LINC Static gels with only acceptor fluorophores does not change over time.

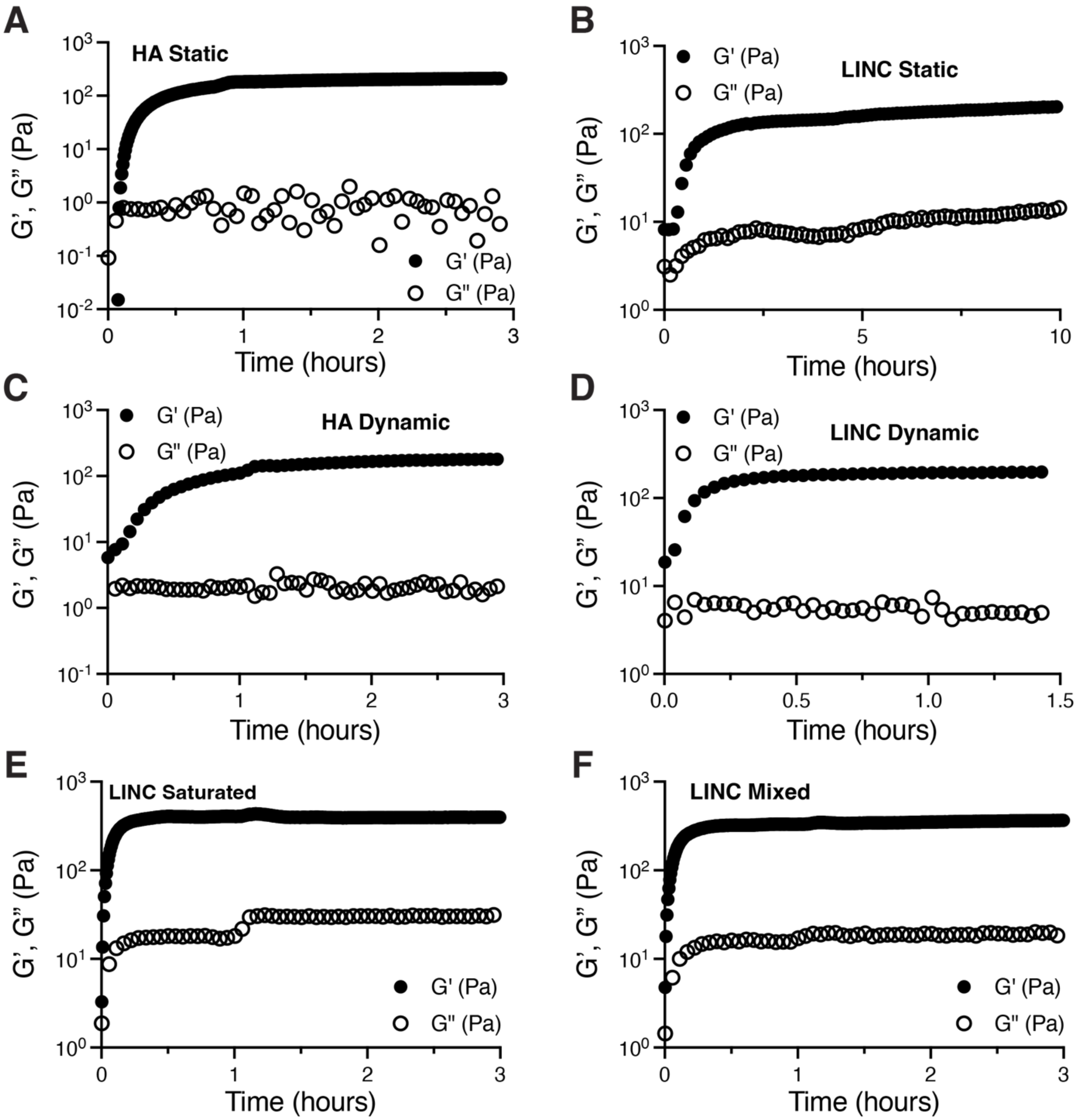


**Fig. S8.** Representative oscillatory rheology time sweeps. **(A)** HA Static, **(B)** LINC Static, **(C)** HA Dynamic, **(D)** LINC Dynamic, **(E)** LINC Saturated, and **(F)** LINC Mixed gels.

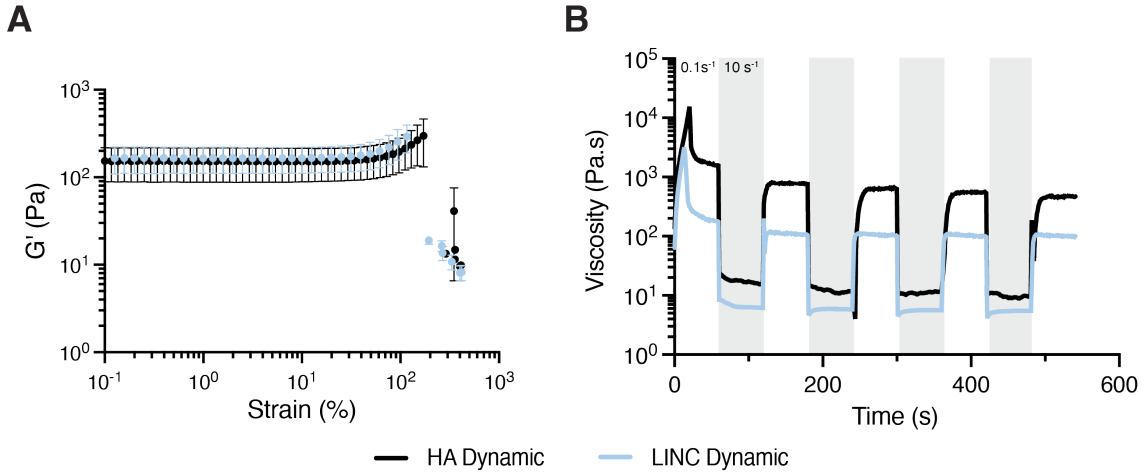


**Fig. S9.** Rheological characterization of strain stiffening, shear-thinning, and recovery. **(A)** Oscillatory strain sweep of HA Dynamic and LINC Dynamic gels showing strain stiffening behavior. N=2, data are mean ± standard deviation. **(B)** Representative viscosity measurements during alternating application of high (10 $s^{-1}$) and low (1 $s^{-1}$) shear rates demonstrating shear-thinning and recovery behavior.

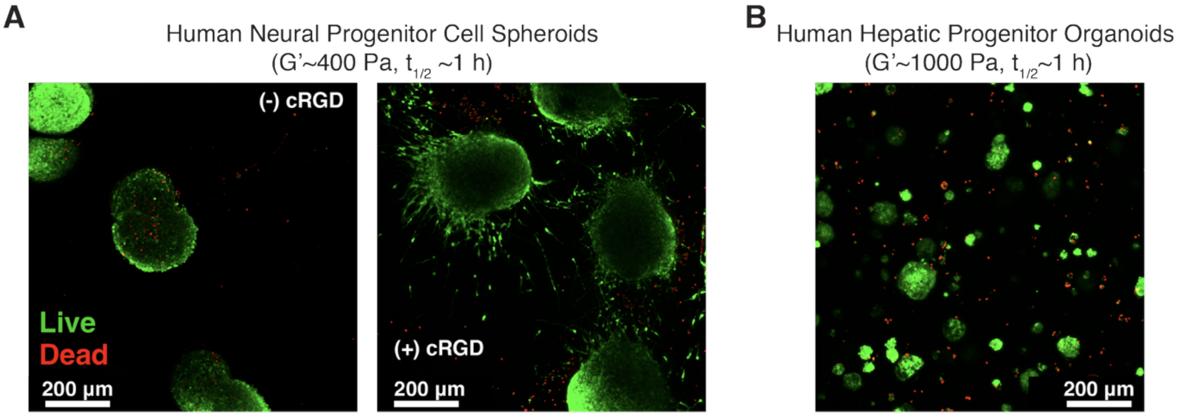


**Fig. S10.** Cell viability staining in LINC gels. Live (calcein AM, green) and dead (ethidium homodimer, red) cells encapsulated within LINC Dynamic gels for 7 days. **(A)** hNPC spheroids with and without cRGD as well as **(B)** human hepatic progenitor organoids with cRGD and laminin.

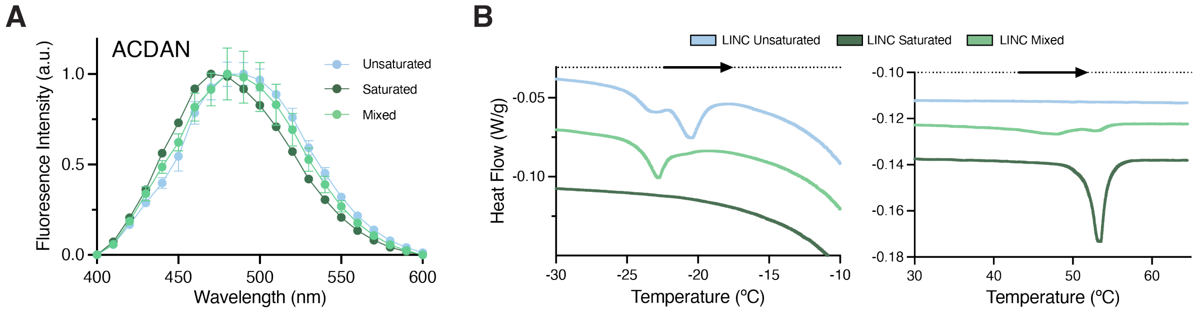


**Fig. S11.** Membrane fluidity and DSC of LINC Mixed. **(A)** ACDAN fluorescence spectra of LINC Unsaturated, Saturated, and Mixed gels. **(B)** DSC heating curves of LINC Unsaturated, Saturated, and Mixed gels.

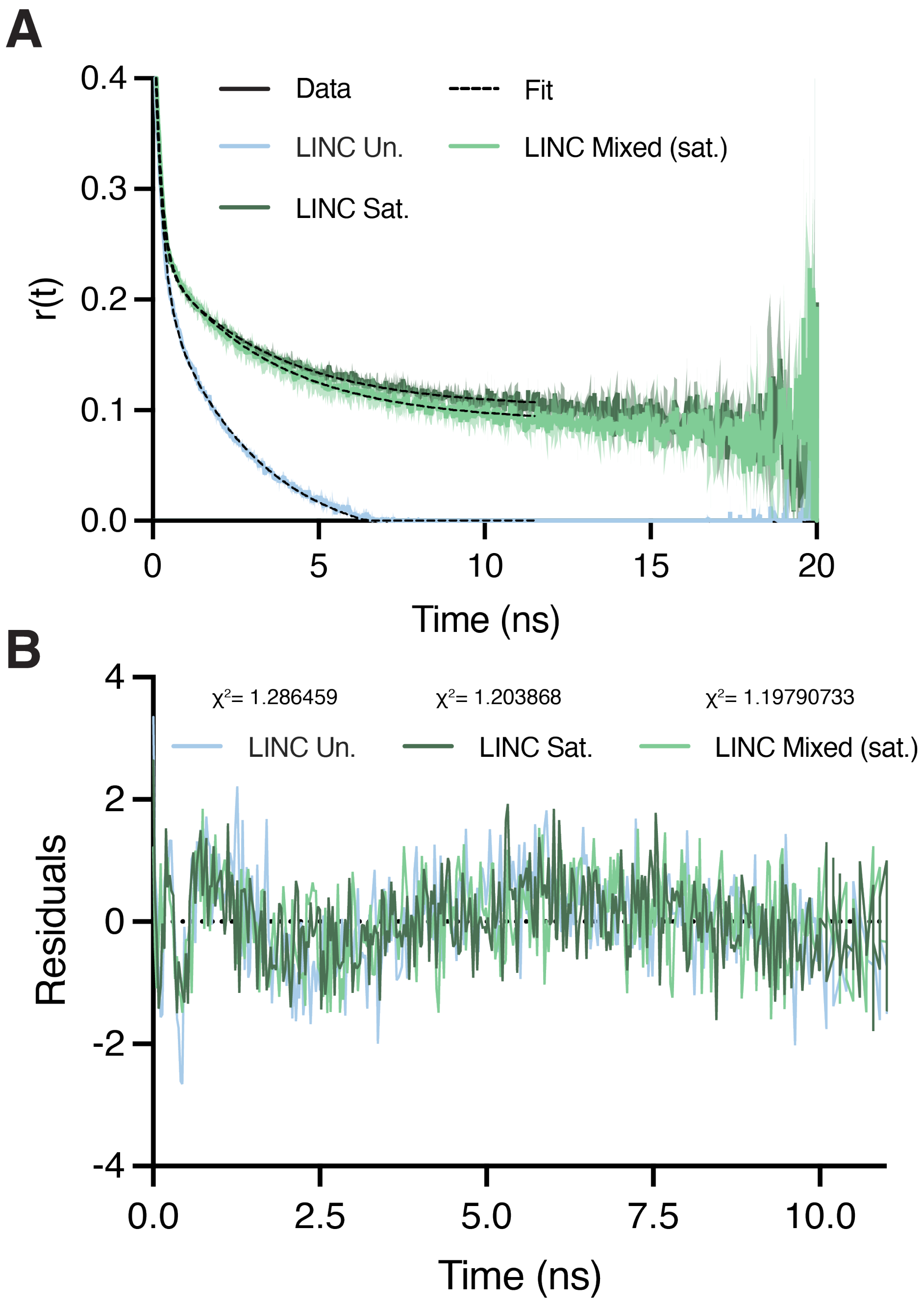


**Fig. S12.** Liposome fluorescence anisotropy. **(A)** Full time correlated fluorescence anisotropy curves and **(B)** fit residuals and $\chi^2$ over the fitted time (~11 ns). Due to the wave-like structure of the residuals, we opted to compare overall anisotropy curve trends rather than extracting fitted parameters to minimize effects of experimental error in our analysis. N=3, data are mean ± standard deviation.

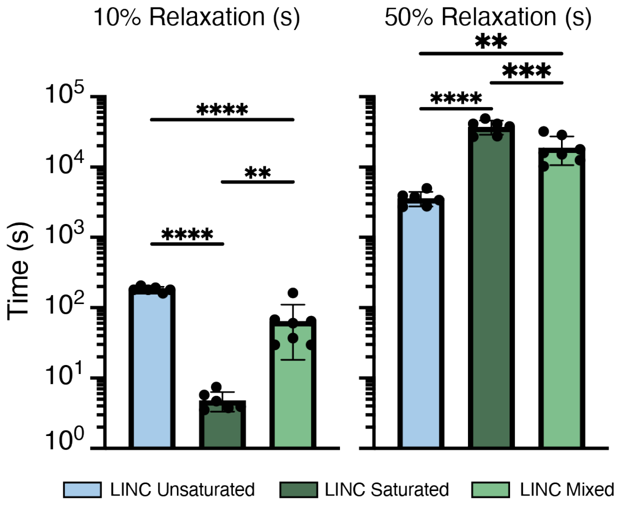


**Fig. S13.** LINC gel stress relaxation. Time required for 10% and 50% stress relaxation in the LINC Unsaturated, Saturated, and Mixed gels. Each plotted point is a single replicate. Data are mean ± standard deviation. 10%: **p=.0061, ****p<0.0001. 50%: **p=0.003, ***p=0.0006, ****p<0.0001. Statistical analysis was one-way ANOVA with Tukey's multiple comparisons test.

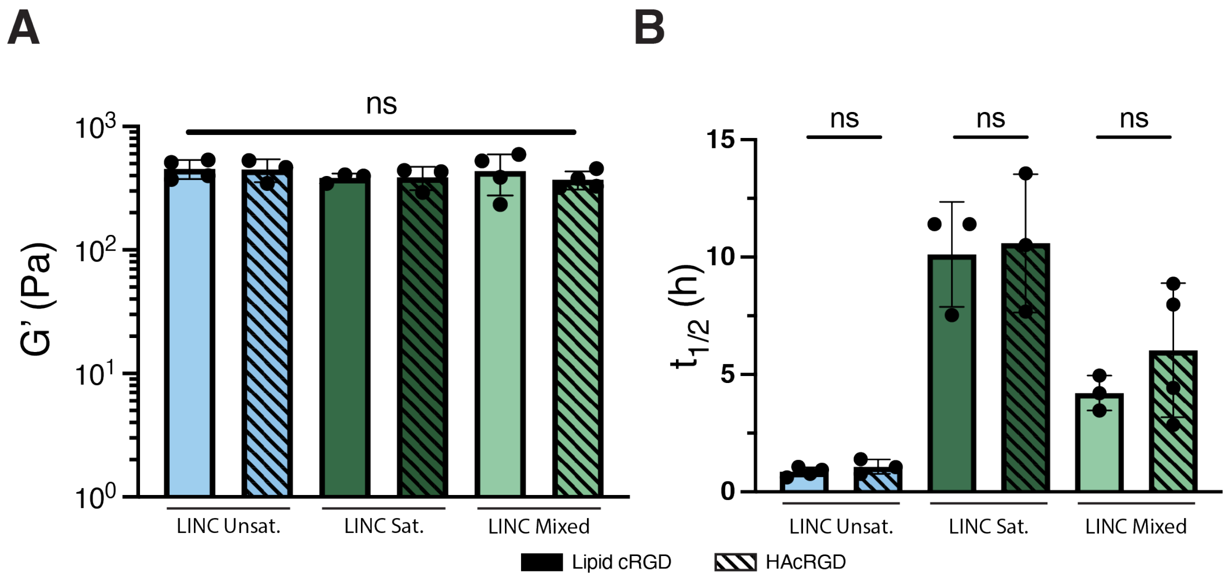


**Fig. S14.** Viscoelasticity of LINC gels with cRGD. Impact of cRGD placement on **(A)** plateau storage modulus and **(B)** stress relaxation $t_{1/2}$ in LINC Unsaturated, Saturated, and Mixed gels. Statistical analysis was one-way ANOVA with Tukey's multiple comparisons test (A) and two-tailed unpaired *t* tests (B). $p > 0.05$ was considered not statistically significant (ns).

**Table S1.** Summary of liposome formulations for LINC gels. Lipid concentration was kept constant in all gels.

| Condition Name | Lipids | Lipid mol Ratio | Representative Diameter (nm) | Representative PDI | Functionalization | Hydrazine:PEG |
|---|---|---|---|---|---|---|
| LINC Static | DOPC/DOPE-mal | 92/8 | 104.8 | 0.267 | Norbornene | - |
| LINC Unsaturated 200 Pa | DOPC/DOPE-mal | 94.25/5.75 | 109.2 | 0.275 | Hydrazine | 63.75:36.25 |
| LINC Unsaturated - 0% PEG | DOPC/DOPE-mal | 94.25/5.75 | 117.4 | 0.26 | Hydrazine | 100:0 |
| LINC Unsaturated - 1.44% PEG | DOPC/DOPE-mal | 94.25/5.75 | 100.8 | 0.276 | Hydrazine | 75:25 |
| LINC Unsaturated - 2.72% PEG | DOPC/DOPE-mal | 92.5/7.5 | 111.4 | 0.245 | Hydrazine | 63.75:36.25 |
| LINC Unsaturated - 4.5% PEG | DOPC/DOPE-mal | 91/9 | 136.4 | 0.247 | Hydrazine | 50:50 |
| LINC Unsaturated - 70 Pa | DOPC/DOPE-mal | 96.5/3.5 | 102.1 | 0.265 | Hydrazine | 63.75:36.25 |
| LINC Unsaturated - 700 Pa | DOPC/DOPE-mal | 83/12 | 106.3 | 0.276 | Hydrazine | 63.75:36.25 |
| LINC Unsaturated- 1000 Pa | DOPC/DOPE-mal | 85/15 | 95.5 | 0.257 | Hydrazine | 63.75:36.25 |
| LINC Unsaturated - 2000 Pa | DOPC/DOPE-mal | 80/20 | 98.8 | 0.258 | Hydrazine | 63.75:36.25 |
| LINC Unsaturated - 600 Pa | DOPC/DOPE-mal | 70/30 | 90.8 | 0.274 | Hydrazine | 63.75:36.25 |
| LINC Unsaturated 400 Pa | DOPC/DOPE-mal | 92.5/7.5 | 111.4 | 0.245 | Hydrazine | 63.75:36.25 |
| LINC Unsaturated 400 Pa RGD | DOPC/DOPE-mal/DOPE-DBCO | 90.5/7.5/2 | 105.3 | 0.258 | Hydrazine | 63.75:36.25 |
| LINC Saturated 400 Pa RGD | DSPC/DSPE-mal/DPPE-DBCO | 92.75/5.25/2 | 100.9 | 0.176 | Hydrazine | 100:0 |
| LINC Phase Separated 400 Pa RGD | DOPC/DSPC/DOPE-mal/DPPE-DBCO | 25.5/65/7.5/2 | 112.8 | 0.232 | Hydrazine | 100:0 |
| LINC Saturated 400 Pa | DSPC/DSPE-mal | 94.75/5.25 | 106.1 | 0.256 | Hydrazine | 100:0 |
| LINC Phase Separated 400 Pa | DOPC/DSPC/DOPE-mal | 25.5/67/7.5 | 107 | 0.231 | Hydrazine | 100:0 |

**Table S2.** Summary of HA characteristics for selected gel conditions. Concentration, functionalization, and molecular weight were held constant across conditions.

| Condition Name | Total Concentration (wt%) | Functionalization | Functionalization (%) | Molecular Weight (kDa) |
|---|---|---|---|---|
| HA Static | 1 | Tetrazine and Norbornene | 8, 32 | 100 |
| HA Dynamic | 1 | Benzaldehyde and Hydrazine | 20, NA | 100 |
| LINC Static | 1 | Tetrazine | 8 | 100 |
| LINC Unsaturated | 1 | Benzaldehyde | 20 | 100 |
| LINC Saturated | 1 | Benzaldehyde | 20 | 100 |
| LINC Mixed | 1 | Benzaldehyde | 20 | 100 |
